# β-catenin perturbations control differentiation programs in mouse embryonic stem cells

**DOI:** 10.1101/2020.05.15.098137

**Authors:** Elisa Pedone, Mario Failli, Gennaro Gambardella, Rossella De Cegli, Antonella La Regina, Diego di Bernardo, Lucia Marucci

**Affiliations:** Department of Engineering Mathematics, University of Bristol, Bristol BS8 1UB, UK; School of Cellular and Molecular Medicine, University of Bristol, Bristol BS8 1TD, UK; Telethon Institute of Genetic and Medicine Via Campi Flegrei 34, 80078 Pozzuoli, Italy; Department of Chemical, Materials and Industrial Production Engineering, University of Naples Federico II, 80125 Naples, Italy; Department of Electrical Engineering and Information Technology, University of Naples Federico II, 80125 Naples, Italy; BrisSynBio, Bristol BS8 1TQ, UK

## Abstract

The Wnt/β-catenin pathway is involved in development, cancer and embryonic stem cell (ESC) maintenance; its dual role in stem cell self-renewal and differentiation is still controversial. Here, by applying an *in vitro* system enabling inducible gene expression control, we report that moderate induction of transcriptionally active exogenous β-catenin in β-catenin null mouse ESCs promotes epiblast-like cell (EpiLC) derivation *in vitro*. Instead, in wild type cells moderate chemical pre-activation of the Wnt/β-catenin pathway promotes EpiLC *in vitro* derivation. Finally, we suggest that moderate β-catenin levels in β-catenin null mouse ESCs favour early stem cell commitment towards mesoderm if the exogenous protein is induced only in the ‘ground state’ of pluripotency condition, or endoderm if the induction is maintained during the differentiation. Overall, our results confirm previous findings about the role of β-catenin in pluripotency and differentiation, while indicating a role for its doses in promoting specific differentiation programs.

## Introduction

Pluripotent Cells (PCs) are characterized by indefinite proliferative and differentiation potential and their identity is determined by the balance between signals promoting self-renewal and differentiation. The first step for stem cell differentiation is the exit from the pluripotent state, tightly controlled by signalling pathway and gene regulatory networks which can drive specific lineage commitment. During murine development *in vivo*, embryonic stem cells (hereafter called ESCs), that represent the naïve pluripotent state of the early epiblast (Evans and Kaufman, 1981, Martin, 1981, Ying et al., 2008), convert into the late epiblast and finally in terminally differentiated somatic cells. ESCs can be derived from the pre-implantation epiblast; they provide an excellent system for understanding signalling pathway interplay in cell fate decision making.

In serum-based cultures, mouse ESCs are heterogeneous for the expression of pluripotency genes (Martin, 1981, Evans and Kaufman, 1981, Brook and Gardner, 1997, Williams et al., 1988, Smith et al., 1988, Ying et al., 2003a, Niwa et al., 1998, Matsuda et al., 1999, Marucci et al., 2014), while, when cultured in serum-free media supplemented with inhibitors of MEK1/2 (PD) and GSK3α/β (Chiron) and in presence or not of the Leukaemia Inhibitory Factor-LIF (2i or 2i+LIF) (Ying et al., 2008), a uniform self-renewal condition known as ‘ground state’ of pluripotency is established; it is characterized by homogenous gene expression (Boroviak et al., 2015, Marks et al., 2012, Godwin et al., 2017, Ghimire et al., 2018), genome demethylation (Ficz et al., 2013, Habibi et al., 2013, Leitch et al., 2013) and naïve pluripotency (Nichols and Smith, 2012, Alexandrova et al., 2016). Following release from 2i or 2i+LIF, epiblast-like cells (hereafter called EpiLCs) appear *in vitro* as an intermediate of ESC differentiation (Chen et al., 2018, Hayashi et al., 2011, Buecker et al., 2014, Krishnakumar et al., 2018). EpiLCs are transcriptionally comparable to epiblast stem cells (hereafter called EpiSCs), although the latter better resemble cells of the anterior primitive steak (Kojima et al., 2014). EpiSCs, derived from the post-implantation epiblast, are capable of differentiating in all the germ-layers (Brons et al., 2007, Tesar et al., 2007); they differ from ESCs in morphology, clonogenicity, gene expression, epigenome status and, most importantly, ability to contribute to chimaeras (Ghimire et al., 2018, Tesar et al., 2007, Brons et al., 2007, Han et al., 2010). EpiSCs require ActivinA and the fibroblast growth factor 2 (FGF2) (Brons et al., 2007, Tesar et al., 2007) for *in vitro* expansion; FGF signalling pathway activation, while promoting EpiSC self-renewal, induces ESC differentiation (Stavridis et al., 2007, Kunath et al., 2007, Guo et al., 2009). Different protocols based on FGF2 treatment, in combination or not with ActivinA and inhibitors of the LIF/STAT3 and the Wnt/β-catenin pathways, have been proposed for the derivation and expansion of EpiLCs and EpiSCs both in serum-based and serum-free culture conditions (Joo et al., 2014, Sumi et al., 2013, Kurek et al., 2015, Hayashi et al., 2011, Gouti et al., 2014, Tsukiyama and Ohinata, 2014). Self-renewing EpiSCs have been established by simultaneous activation and inhibition of the Wnt/β-catenin pathway (Kim et al., 2013); however, the effect of these perturbations on the ESCs-EpiLCs transition has not been fully explored.

The Wnt/β-catenin is a highly conserved signalling pathway involved in ESCs self-renewal (Sato et al., 2004) and cell-cycle progression (De Jaime-Soguero et al., 2017). β-catenin levels are tightly controlled by the active transcription of negative regulators working at different levels of the signalling cascade (Stamos and Weis, 2013): Axin2 (Jho et al., 2002, Chia and Costantini, 2005, Leung et al., 2002) is part of the destruction complex whereas DKK1 (Glinka et al., 1998) binds to the Wnt receptor complex attenuating cellular response upon activation of the pathway. These negative feedback loops contribute to the emergence of nonlinear dynamics in the Wnt/β- catenin pathway, proved to be important in different biological and developmental aspects (see (Pedone and Marucci, 2019) for a review), ESCs pluripotency and somatic cell reprogramming (Marucci et al., 2014, Aulicino et al., 2014, Ho et al., 2013, Kimura et al., 2016, Lluis et al., 2008).

The role of the canonical Wnt pathway in early *in vivo* developmental stages and the requirement of its activation for ESC self-renewal have been a matter of intense research (Anton et al., 2007, Ying et al., 2008, Soncin et al., 2009, Wagner et al., 2010, Lyashenko et al., 2011, Wray et al., 2011, Faunes et al., 2013, Ye et al., 2017, Chatterjee et al., 2015, Ortmann et al., 2020, Tao et al., 2020, Theka et al., 2019, Aulicino et al., 2020). Pluripotency incompetence of β-catenin^-/-^ ECSs has been reported in two independent studies (Anton et al., 2007, Wagner et al., 2010); this phenotype was contradicted later using newly generated β-catenin^-/-^ cell lines, which showed self-renewal in both serum and 2i+LIF (hereafter called 2i/L), but presented some differentiation defects when LIF-deprived (Wray et al., 2011, Lyashenko et al., 2011, Aulicino et al., 2020). Such knock-out models provide an excellent *in vitro* system to study β-catenin function on ESC decision making.

Here, we take advantage of the β-catenin^-/-^ ESC line generated by Aulicino and colleagues (Aulicino et al., 2020), where the entire β-catenin coding sequence was removed to avoid possible compensatory mechanisms from aberrant truncated isoforms, to study the effect of β-catenin perturbations on the exit from pluripotency and differentiation. Different β-catenin doses have been indirectly achieved in the past by mutating the adenomatous polyposis coli gene (APC) (Kielman et al., 2002); teratomas from the mutants with the highest β-catenin transcriptional activity showed major differentiation defects in the neuroectoderm, dorsal mesoderm and endoderm lineages. Of note, results in (Kielman et al., 2002) suggest that active β-catenin nuclear translocation (different across mutants) might also be involved in the observed differentiation impairment.

Models enabling direct modulation of β-catenin can be used to systematically associate protein perturbations to pluripotency and differentiation phenotypes. For this aim, we tuned β-catenin levels in β-catenin^-/-^ ESCs applying an improved inducible system (Pedone et al., 2019) and measured both global gene expression in ground state pluripotency (i.e., 2i/L) and following 2i/L withdrawal, as well as the efficiency of ESC-EpiLC transition *in vitro*. We demonstrated that moderate β-catenin activation in β-catenin^-/-^ ESCs (between null and wild type levels) and moderate chemical pre-activation of the Wnt/β-catenin pathway in wild type ESCs promote efficient EpiLC *in vitro* derivation. Finally, the transcriptome of β-catenin^-/-^ ESCs expressing different doses of exogenous β-catenin before and/or during differentiation confirmed what we and others reported about the dispensable requirement of β-catenin transcriptional activity for pluripotency establishment (Lyashenko et al., 2011, Pedone et al., 2019, Wray et al., 2011, Faunes et al., 2013, Aulicino et al., 2020), while suggesting that specific β-catenin perturbations cause a bias towards the endoderm lineage, in line with Lef-1 related results previously reported (Ye et al., 2017).

Overall, our study highlights that a synergistic effect of β-catenin doses and culture conditions controls *in vitro* ESC fate decision making at the exit from pluripotency.

## Results

### Wnt/β-catenin pathway perturbations control *in vitro* generation of EpiLC

To study the role of the Wnt/β-catenin pathway in EpiLC *in vitro* derivation, we used the C1-EF1a-rtTA_TRE3G-DDmCherryβ-catenin^S33Y^ (hereafter called C1) ESC line we previously generated (Pedone et al., 2019). Briefly, β-catenin^−/−^ ESCs (Aulicino et al., 2020) were modified to stably express a doxycycline-inducible fusion protein comprising the conditional destabilising domain (DD), the mCherry fluorescent protein and the constitutive active β-caten in^S33Y^ (Figure 1A) (Sadot et al., 2002). The inducer molecule doxycycline (dox) enables transcriptional initiation, while trimethoprim (TMP) allows protein stabilisation by inactivating the DD (Figure 1A) (Pedone et al., 2019). The use of a constitutively active and conditional β-catenin^S33Y^ form, uncoupled from upstream endogenous regulations and in a knock-out background, avoids possible issues (i.e., compensatory mechanisms, genetic variation, off-target effects) resulting from endogenous protein induction or chemical pathway activation.

**Figure 1.**
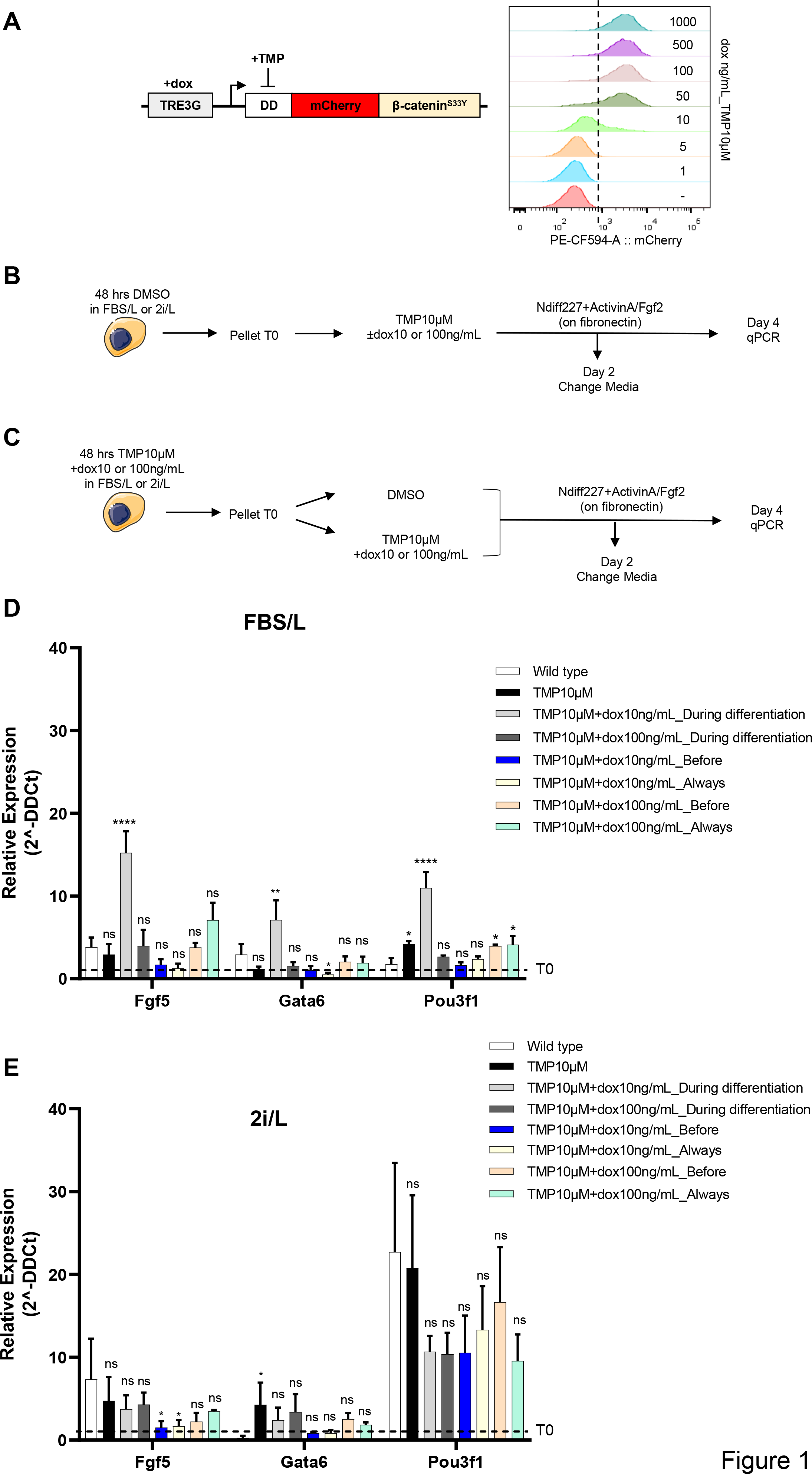
Dual-input control of β-catenin doses in EpiLC *in vitro* derivation (β- catenin^-/-^ background). **A** Dual-input regulation system consisting of the doxycycline responsive element and the conditionally destabilised mCherryβ-catenin^S33Y^ module. Doxycycline (dox) and trimethoprim (TMP) allow mCherryβ-catenin^S33Y^ transcription initiation and protein stabilisation, respectively. **(A, inset)** Flow cytometry profile of C1 ESCs treated for 24 hrs with TMP10µM and the indicated concentrations of dox. **B, C** Experimental scheme ESC to EpiLC conversion. C1 ESCs cultured in FBS/L or 2i/L, were pre-treated either with DMSO **(B)** or TMP10µM and dox10-100ng/mL **(C)**. Following 48 hrs of treatment, cells were seeded on fibronectin in NDiff227 and exposed to ActvinA/FGF2 and different combinations of DMSO, dox and TMP for 4 days before being collected for RNA extraction. After 2 days, the media was changed, and the drugs were refreshed. **D, E** Fgf5, Gata6 and Pou3f1 expression in C1 ESCs cultured in FBS/L **(D)** or 2i/L (**E)** and differentiated for 4 days in NDiff227+ActivinA/FGF2 and different combination of DMSO, doxy and TMP. Data are represented as fold-change with respect to the corresponding pluripotent condition (i.e., time zero before differentiation (T0)) indicated with a dashed line. Data are means± SEM (n=3 biological replicates). p-values from one-way ANOVA with Bonferroni’s multiple comparison test computed over the wild type ESCs are shown, *p<0.05, **p<0.01, ***p<0.001, ****p<0.0001.

We confirmed in C1 cells the correct induction (Figure 1A, inset and (Pedone et al., 2019)), intracellular distribution and functionality of the exogenous protein upon input administration (Figure S1A and (Pedone et al., 2019)). We found a dose-dependent upregulation of the β-catenin target gene Axin2 in C1 ESCs cultured with dox and TMP for 48 hrs, although not reaching activation comparable to Chiron 3µM-treated wild type cells in the case of maximum induction (Figure S1A). This result confirms our previous measurements of the total exogenous β-catenin levels induced by drugs being lower than wild type condition, while still being active in the nucleus, thanks to the use of a mutant form, insensitive to the endogenous degradation machinery (Pedone et al., 2019).

The β-catenin^−/−^ cell line we used had already been characterised for having a transcriptional profile similar to wild type cells in pluripotent (serum/LIF) cultures, with Wnt signalling activation repressing ESC spontaneous differentiation in dependence of β-catenin (Aulicino et al, 2020). We also previously confirmed the dispensable role of β-catenin in pluripotent culture conditions and showed, using Alkaline Phosphatase (AP) staining, that moderate β-catenin induction with our inducible system (using TMP10µM_dox10ng/mL) can protect cells from exiting pluripotency in the absence of both serum and LIF (Pedone et al., 2019).

Following these results, we measured the efficiency of EpiLCs derivation when different doses of exogenous β-catenin are induced in β-catenin^−/−^ cells under pluripotent conditions and/or during differentiation. To appreciate cellular response changes depending on the culture condition, C1 ESCs were expanded either in serum/LIF (hereafter called FBS/L) or in 2i/L. As prolonged culture in 2i/L results in epigenetic changes impairing normal differentiation *in vitro* and development *in vivo* (Choi et al., 2017), we opted for a short-term culture in 2i/L (3 passages).

ESCs from FBS/L or 2i/L (Figures 1B and 1C) were cultured for 48 hrs either in DMSO (Figure 1B) or in the presence of maximum TMP (10µM) combined with low (10ng/mL) or saturating (100ng/mL) dox (Figure 1C). The concentrations of dox were extrapolated from flow cytometry measurements of the mCherry signal to provide two doses (moderate and high) of the exogenous protein (Figure 1A, inset). To explore the effect of β-catenin perturbations on ESC-EpiLC transition, we adapted the protocol for EpiSCs culture from (Kim et al., 2013) (see STAR Methods for details), FBS/L- or 2i/L-cultured C1 ESCs were supplemented with ActivinA, FGF2 (Kim et al., 2013) and different combinations of DMSO, TMP and dox (Figures 1B, 1C and STAR Methods). Cells were kept under these conditions for 4 days, with media refreshed after the first 2 culture days; flow cytometry showed that the mCherry florescence was only marginally influenced by the frequency media and drugs are refreshed (Figure S1B). The fluorescent reporter was expressed in a dose-dependent manner following 48 hrs drug treatment in pluripotency conditions (Figures S1C and S1D); similarly, cells were sensitive to different concentrations of drugs, and their removal, during differentiation (Figures S1E and S1F). Upon protocol completion, ESCs were analysed for the expression of the early epiblast and epiblast-like markers Fgf5, Gata6 and Pou3f1 (i.e., Oct6) (Kalkan et al., 2017, Navarra et al., 2016, Wang et al., 2021), and of the pluripotency markers Nanog and Esrrb (Kalkan et al., 2017) by qPCR (Figures 1B-1E, S1G and S1H). TMP-treated (i.e., control) C1 cells did not show a significant differentiation impairment, as compared to the wild type parental cell line (Figures 1D and 1E). We found a significant upregulation of epiblast-like genes in FBS/L cultured C1 ESCs induced with a low amount of dox (TMP10µM_dox10ng/mL “During Differentiation” sample, Figure 1D), suggesting that this culture condition favours EpiLC conversion (Figure 1D). This effect was not observed in C1 ESCs pre-cultured in 2i/L for 3 passages (Figure 1E). As expected, upon the differentiation process, Nanog and Esrrb were efficiently downregulated (Figures S1G and S1H).

Altogether, these results indicate that both culture media and β-catenin doses strongly influence how cells respond to the ActivinA/FGF2 stimulus, with moderate β-catenin induction during the differentiation protocol facilitating the transition towards EpiLCs of FBS/L-cultured ESCs.

### Chemical activation of the canonical Wnt pathway in pluripotent conditions modulates EpiLC conversion of wild type ESCs

The above results motivated us to explore the EpiLC conversion potential of wild type ESCs when deprived of pluripotency factors and exposed to different chemical perturbations of the endogenous Wnt/β-catenin pathway. Simultaneous activation/inhibition of the canonical WNT pathway has been previously reported to facilitate EpiSCs derivation and *in vitro* expansion (Kim et al., 2013). However, a potential effect of a such combination was not explored in the ESC-EpiLC conversion. When activating the pathway with the Gsk3 inhibitor Chiron-99021 (Chiron), the levels of activated β-catenin will be here significantly higher than in the experiments in Figure 1, thus different results in EpiLC conversion are expected.

We measured the transition from ESCs to EpiLCs in a 4-day time-course by qPCR (Figure 2A). Before differentiation, ESCs were cultured in FBS/L and treated for 48 hrs with DMSO or with Chiron to pre-activate the canonical Wnt pathway, or were maintained in 2i/L for 3 passages (i.e., 1 week; Figure 2A). At Day 0, cells were exposed to different combination of drugs added to the NDiff227 (Guo et al., 2009): ActivinA+FGF2+DMSO (AFD); ActivinA+FGF2+Ch1µM (AFCh1); ActivinA+FGF2+Ch3µM (AFCh3); Ch1µM (Ch1); Ch1µM+XAV2µM (Ch1X2); Ch3µM (Ch3); Ch3µM+XAV2µM (Ch3X2) (Kim et al., 2013) (Figure 2A). The expression of the epiblast-like genes Fgf5, Gata6 and Pou3f1 and the pluripotency genes Nanog and Esrrb was analyzed by qPCR after 4 days. A change of media was performed after the first 2 culture days (Figure 2A).

**Figure 2.**
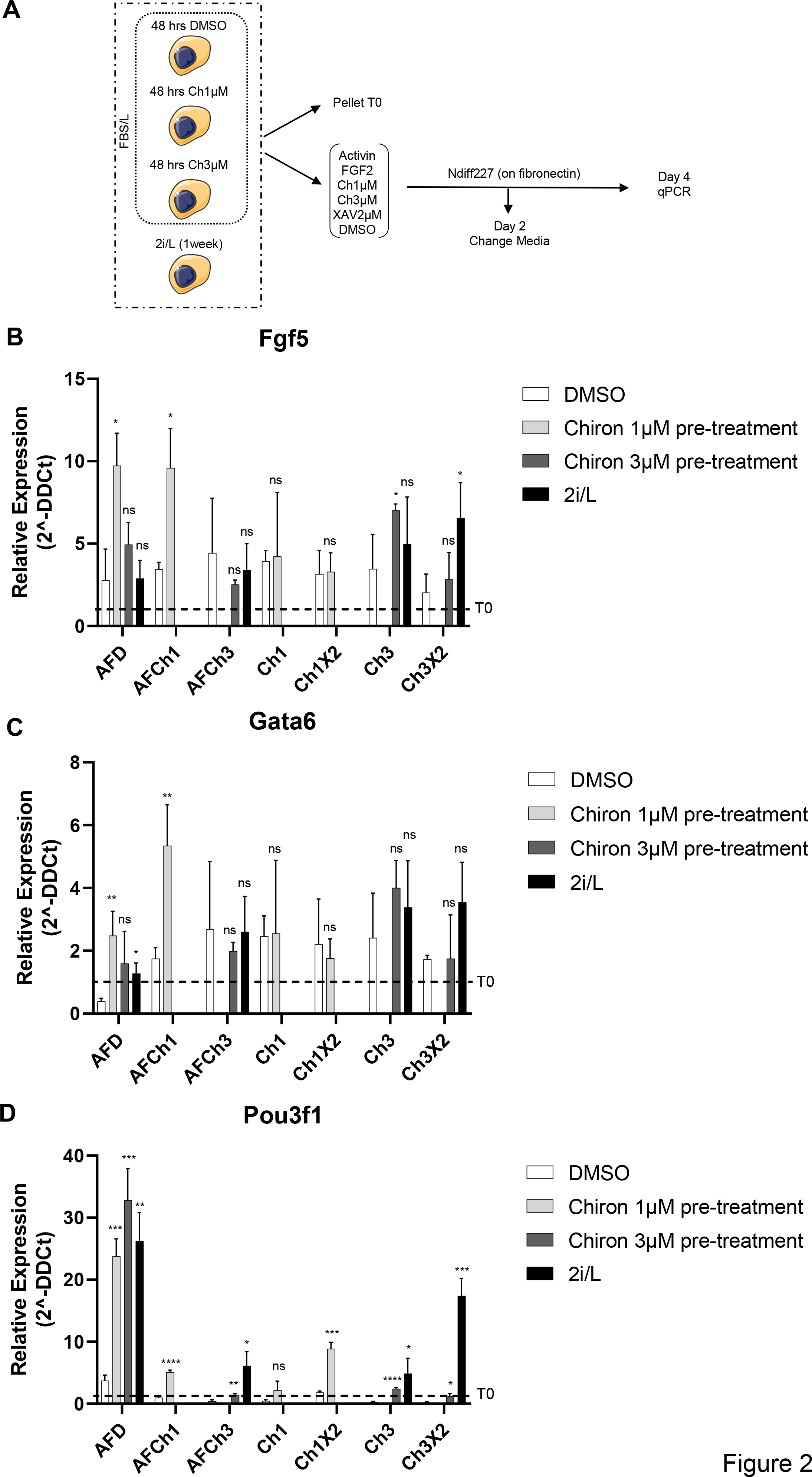
Chemical perturbation of the Wnt/β-catenin in EpiLC *in vitro* derivation (β-catenin wild type background). **A** Experimental scheme of EpiLCs derivation. Wild type ESCs cultured in FBS/L and pre-treated for 48 hrs with DMSO and Chiron (1-3 µM), or in 2i/L for 3 passages, were seeded on fibronectin in NDiff227 supplemented with different combinations of drugs (ActivinA+FGF2+DMSO (AFD); ActivinA+FGF2+Ch1µM (AFCh1); ActivinA+FGF2+Ch3µM (AFCh3); Ch1µM (Ch1); Ch1µM+XAV2µM (Ch1X2); Ch3µM (Ch3); Ch3µM+XAV2µM (Ch3X2)). After 2 days, the media was changed, and the drugs were refreshed. Expression of epiblast-like genes was measured by qPCR in pluripotent conditions (T0) and after 4 days of differentiation. **B-D**Fgf5 **(B)**, Gata6 **(C)** and Pou3f1 **(D)** expression in in DMSO, Ch1µM, Ch3µM and 2i/L pre-cultured wild type ESCs differentiated for 4 days in NDiff227 and the indicated combination of drugs. Data are represented as fold-change with respect to the corresponding pluripotent condition (i.e., time zero before differentiation (T0)) indicated with a dashed line. Data are means± SEM (n=3 biological replicates). p-values from two-tailed unpaired t-test computed over the DMSO are shown, *p<0.05, **p<0.01, ***p<0.001, ****p<0.0001.

In the FBS/L condition, the expression of epiblast-like genes, as compared to the standard differentiation protocol based on ActivinA and FGF2 (i.e., AFD), showed mixed behaviours upon Wnt/β-catenin pathway perturbations (Figure S2A); however, pluripotency genes were significantly upregulated *vs* AFD in the majority of perturbations (Figure S2E), suggesting that the standard differentiation protocol, with no activation/inhibition of the pathway, is the most suited to support the ESC-EpiLC transition in FBS/L.

Next, we differentiated cells after pre-activation of the Wnt/β-catenin pathway in FBS/L condition (Figures S2B, S2C, S2F and S2G) or 1 week of pre-culture in 2i/L (Figures S2D and S2H). Upon Chiron 1µM, but not Chiron 3 µM, pre-treatment in FBS/L, the expression of EpiLC genes with the AFD protocol was higher as compared to the DMSO condition (Figures 2B-2D and S2A-S2C). In Chiron 1µM pre-treated and AFD- differentiated ESCs, the downregulation of pluripotency genes was comparable to that observed in 2i/L cultured cells, which can efficiently differentiate (Figures S2I and S2J, (Hackett et al., 2018, Hayashi et al., 2011)). Instead, pluripotency genes were not efficiently downregulated when the Wnt/β-catenin pathway was perturbed also during differentiation in ESCs pre-activated with Chiron 1-3µM in FBS/L (AFCh1 and Ch1 in Figures S2F and S2J; AFCh3 and Ch3 in Figures S2G and S2I).

Altogether, these results indicate that intermediate chemical pre-activation of the Wnt pathway in FBS/L cultured wild type cells favours the ESC-EpiLC transition using the standard differentiation protocol (AFD).

### Transcriptome and WGCNA analysis of ESC exit from pluripotency with varying β-catenin doses

Next, we studied C1 ESCs exit from the ‘ground state’ of pluripotency, i.e., upon 2i/L withdrawal, by RNA-sequencing; such monolayer differentiation protocol does not induce a specific cell fate and is well suited to observe possible β-catenin dependent differentiation bias (Kalkan et al., 2017).

C1 ESCs cultured in 2i/L for 3 passages (i.e., 1 week) were treated for 48 hrs with saturating concentrations of dox and TMP (100ng/mL and 10µM, respectively) to induce the expression of the exogenous fusion protein (Figure 3A). Taking advantage of the mCherry-tag for exogenous β-catenin induction, dox/TMP-treated C1 ESCs were sorted into two different subpopulations: C1 with Middle and High β-catenin levels (hereafter called C1M and C1H samples, respectively; Figures 3A and S3A). We also checked activation of the pathway in sorted cells by measuring Axin2 via qPCR (Figure S3B) and found a dose-dependent activation of this target gene. Sorted cells were cultured in NDiff227 media± inducers for 4 days before being transcriptionally profiled (see STAR Methods for details; Figure 3A). Sequencing informed about the transcriptome of pluripotent C1, C1M and C1H ESCs, and of their differentiated counterparts (Day 4 samples) cultured in NDiff227 and DMSO (i.e., upon dox/TMP withdrawal; hereafter called C1MV and C1HV), or in NDiff227 and TMP/dox (hereafter called C1MDT and C1HDT) during differentiation. The C1 and C1T samples refer to cells treated only with TMP10µM in the pluripotent and differentiated states, respectively. We measured mCherry levels upon addition/removal of dox and TMP and confirmed dose-/administration time-dependent upregulation of exogenous β- catenin (C1MDT and C1HDT; Figure S3C).

**Figure 3.**
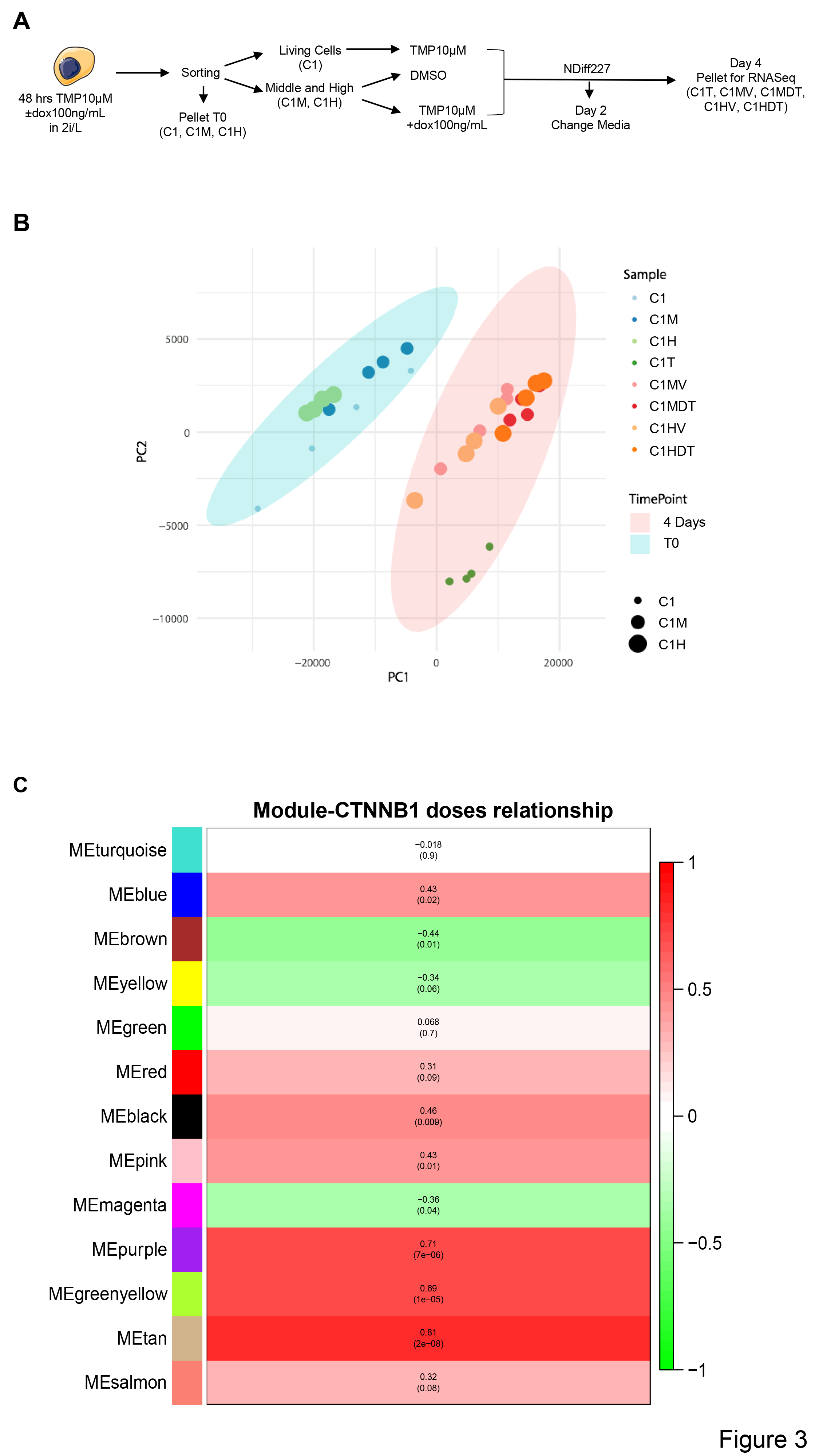
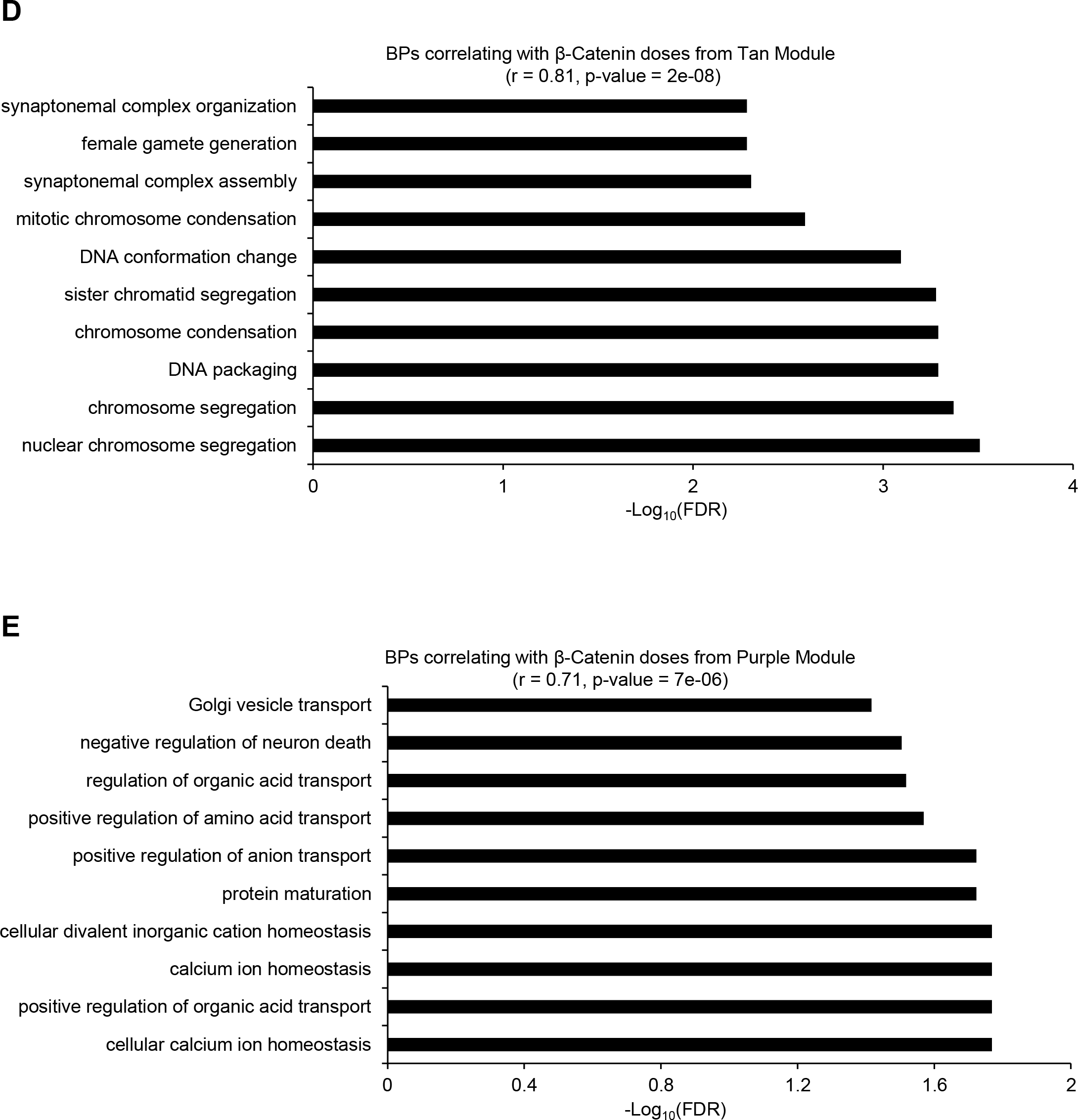
Transcriptome analysis of monolayer differentiation experiments upon β-catenin perturbations and WGCNA of the genes correlating with β-catenin doses. **A** Experimental scheme of the monolayer differentiation protocol. 2i/L C1 ESCs were pre-treated with TMP10µM and dox100ng/mL for 48 hrs; living cells were then sorted from the Dapi negative fraction of TMP-treated cells (C1), whereas β-catenin induced cells from dox/TMP-treated samples were FACS-sorted from the mCherry fraction and divided into Middle (C1M) and High (C1H) subpopulations. 1.5×10^4^ cells/cm^2^ from each individual population were then seeded on gelatin in NDiff227 supplemented with DMSO or TMP±dox100ng/mL. After 4 days of differentiation in NDiff227, cells were collected and processed for the RNA sequencing. During the protocol, the media was changed, and the drugs refreshed after 2 culture days. **B** Principal Component Analysis (PCA) of all samples following batch correction with Combat-seq method; the average of replica is shown. **C** Eigenmodules correlating with β-catenin doses; the Pearson Correlation Coefficient (*r)* and relative p-value are shown. **D, E** Bar-chart of the top-ten enriched biological processes (BP) with FDR < 0.05 from genes belonging to the Tan **(D)** and Purple **(E)** WGCNA modules.

To investigate the extent of the batch effect, we applied the ComBat-Seq method (Zhang et al., 2020) considering sample replicates as different batches and preserving differences among sample types. ComBat-Seq applies a set of statistical corrections to remove the batch effect in the dataset and thus reduce spurious correlations between genes. We then compared, for each sample, the 2D-correlation value between the normalized expression profiles with and without using the ComBat-Seq method (Figure S3D). It can be appreciated that the samples are very similar (i.e., correlations close to 1 for each sample) before and after the batch correction thus suggesting that ComBat-Seq applies a negligible correction. Indeed, the Principal Component Analysis (PCA) after batch correction (Figure 3B) does not substantially differ from the one in Figure S3E (no correction).

Principal Components Analysis (PCA, Figure 3B) showed three main clusters: one group included C1, C1M and C1H in the ‘ground state’ of pluripotency, another group included differentiated C1T ESCs and the final group contained all perturbed samples (C1MV, C1HV, C1MDT and C1HDT) after 4 days of differentiation.

Next, to explore the biological processes associated with β-catenin perturbations, we used Weighted Gene Correlation Network Analysis (WGCNA) (Langfelder and Horvath, 2008), a gene network approach that starting form transcriptional data allows to identify highly co-expressed group of genes (a.k.a. modules) and associate them to the phenotypes or experimental conditions under investigation. By applying WGCNA on our transcriptional data, we identified 13 unique gene modules (Figure S3F; see STAR Methods for details), of which seven (i.e., Green, Blue, Black, Brown, Turquoise, Yellow and Pink; Figure S3G and Table S1) were significantly correlated with differentiation time, thus containing genes playing a key role during differentiation (Figure S3G and STAR Methods). In particular, while four out of the seven gene modules (i.e., Green, Blue, Black and Brown) were positively correlated with differentiation time and thus highly co-expressed (i.e., active) after 4 days of differentiation, three modules (i.e., Turquoise, Yellow and Pink) resulted instead anti-correlated with differentiation time, meaning that those genes are highly co-expressed at T0 but not after 4 days of differentiation. Gene Ontology Enrichment Analysis (GOEA) of the most representative genes (Table S1 and STAR Methods) from the modules highly co-expressed at T0 showed a significant enrichment (FDR < 0.05) in biological processes related to regulation of tissue remodelling, embryonic and forebrain development, stem cell population maintenance, cell homeostasis (Figure S3H and Table S1, Turquoise), nuclear division, meiosis and organelle fission (Figure S3I and Table S1, Pink). On the other hand, gene modules positively correlated with differentiation time (i.e., genes active at day 4) showed a significant enrichment (FDR < 0.05) in biological processes related to translation, rRNA processing, ribosomal biogenesis (Figures S3J, S3L and Table S1, Green and Brown), protein transport, processes associated with cellular respiration (Figure S3K and Table S1, Blue), positive regulation of growth, ncRNA processing and neuronal tube formation and development (Figure S3L and Table S1, Brown). Of note, there were no significantly enriched BPs for the genes from the Black, Yellow and Pink modules (Table S1).

Next, to gain more information about the effects of β-catenin at the exit from pluripotency, we also correlated each module with β-catenin doses (basal, moderate and high, Figure 3C) and found three gene modules gradually increasing their co-expression with increasing β-catenin concentration (i.e., Tan, Purple and Green/Yellow; Figure 3C and Table S1). The biological processes (Table S1 and STAR Methods) corresponding to these modules showed enrichment in cell division (Figure 3D and Table S1, Tan), metabolism and negative regulation of neuronal death (Figure 3E and Table S1, Purple).

Overall, the WGCNA analysis confirmed the expected major transcriptional and metabolic changes associated with the exit from the pluripotent status and confirmed previously reported β-catenin functions on cell survival and proliferation (De Jaime-Soguero et al., 2017).

Next, we performed differential gene expression (DGE) analysis between specific pairs of samples and performed GOEA to identify the involved biological processes (red bars in Figures 4A-4D, S4A and S4B). Starting from the pluripotent condition, when comparing C1M and C1H with C1 we found that the first 10 biological processes with an FDR < 0.05 were mainly related to cell cycle and metabolism (Figures S4A, S4B, Tables S2 and S3). Interestingly, the genes exclusively upregulated in C1H were related to tissue differentiation (i.e., eye morphogenesis and urogenital system development) and DNA methylation involved in gamete generation (Figure S4B and Table S3). Only a few signalling pathways were enriched in C1M ESCs compared to the control cell line C1 (Tables S2 and S3). These results, together with the PCA in Figure 3B, confirm previous observations about the dispensable function β-catenin has in pluripotent culture conditions (Lyashenko et al., 2011, Pedone et al., 2019, Wray et al., 2011, Aulicino et al., 2020) and suggest a bias towards differentiation in C1H ESCs (Figure S4B and Table S3).

**Figure 4.**
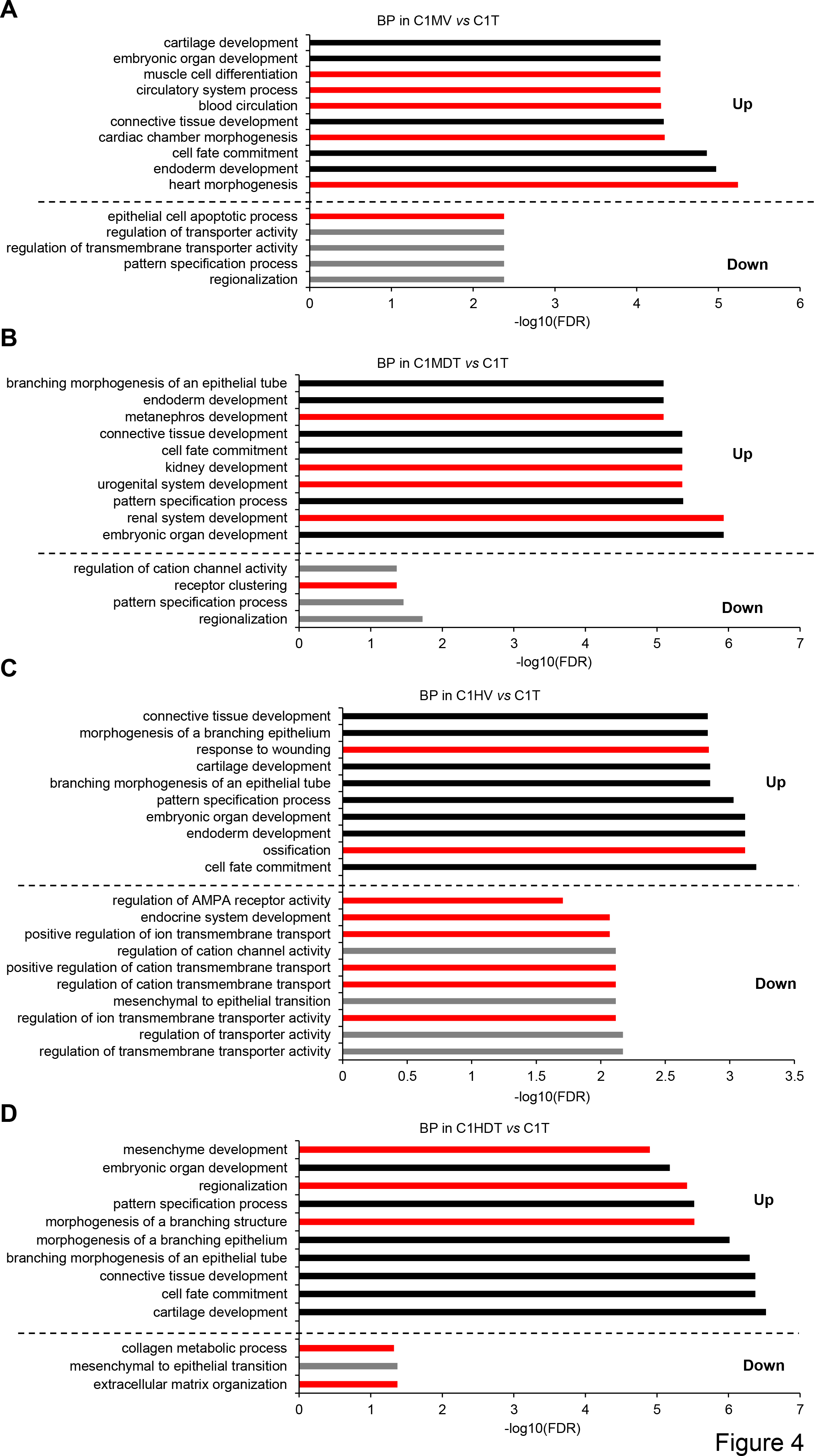
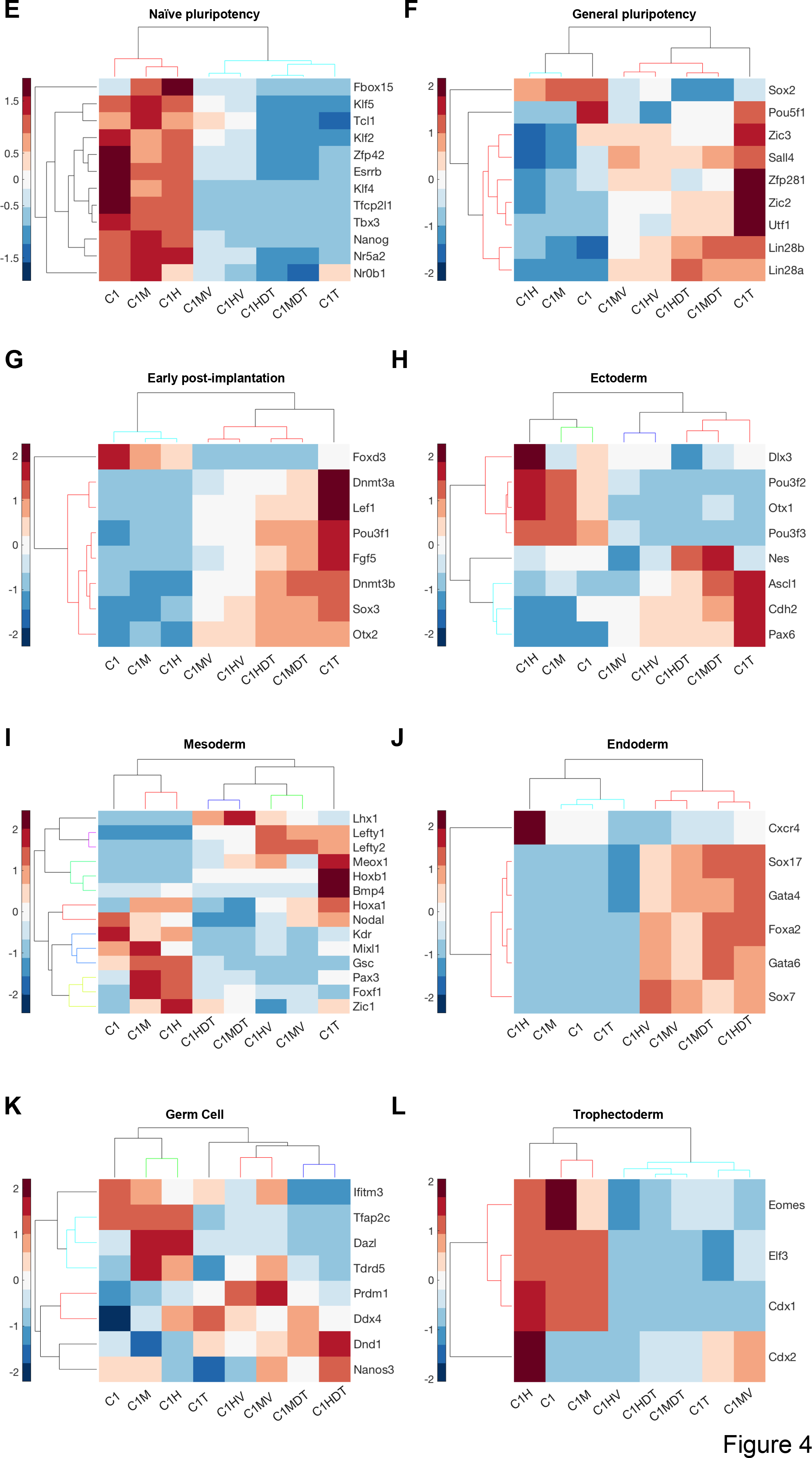
Gene ontology and clustergram of the differential expressed genes in control and perturbed ESCs. **A-D** Bar-chart of the top-ten enriched biological processes (BP) with FDR < 0.05 from differentially expressed genes in C1MV **(A),** C1MDT **(B),** C1HV **(C)** and C1HDT **(D)** compared to C1T ESCs. Black and grey bars represent upregulated and downregulated BPs, respectively. In red bars, the BPs exclusively enriched in the indicated condition. **E-L** Clustergram over heatmaps of Naïve **(E)** and general pluripotency **(H)**, early post-implantation **(G)**, ectoderm **(H)**, mesoderm **(I)**, endoderm **(J)**, germ cell **(K)** and trophectoderm **(L)** lineages from pluripotent and differentiated ESCs expressing different β-catenin amount. Each column is the average of 4 samples from the same experiment.

We then analysed the genes differentially expressed at Day 4 upon β-catenin perturbation as compared to C1T ESCs. The upregulated genes gave the major contribution to the significantly enriched BPs (Figures 4A-4D and Tables S4-S7), while the downregulated genes only contributed to enrich a few processes, namely general metabolic processes (e.g., regulation of transporter and cation channel activity) and mesenchymal to epithelial transition (Figures 4A-4D and Tables S4-S7). Genes exclusively upregulated in the C1MV *vs* C1T comparison belonged to the mesoderm lineage (i.e., cardiovascular system development; Figure 4A and Table S4), while, in the C1MDT *vs* C1T comparison, upregulated genes were enriched for the endoderm lineage (i.e., urogenital system; Figure 4B and Table S5). Nevertheless, mesoderm and endoderm lineages were represented in both comparisons. GO performed on the C1HV and C1HDT comparisons with C1T showed only a few differences in the enriched BPs, that indeed did not define a bias toward a specific lineage (Figures 4C, 4D, Tables S6 and S7).

These results suggest that the major changes in the differentiation program initiated upon 2i/L withdrawal are induced by moderate β-catenin doses and are influenced by the timing of protein induction. The pathway enrichment analysis showed the upregulation of protein metabolism in C1MV and C1HV (Tables S4 and S6), MAPK signalling pathway (Table S4) in C1MV, and ECM-receptor interaction and PI3K-AKT signalling pathway in C1HV (Table S6).

To gauge insights into specific differentiation programs, we selected sets of markers for naïve and general pluripotency, early post-implantation epiblast, ectoderm, mesoderm, endoderm, germ cell and trophectoderm (Kalkan et al., 2017), and clustered our samples according to their expression.

Naïve pluripotency genes were downregulated upon differentiation in all samples, indicating the successful exit of cells from pluripotency (Figure 4E). Pluripotent C1M and C1H samples clustered together (Figure 4E) although close to C1 ESCs confirming that β-catenin is dispensable for pluripotency maintenance. ESCs differentiated in presence of DMSO (i.e., C1MV and C1HV; Figure 4E) clustered together, similarly to samples differentiated in presence of dox and TMP (i.e., C1MDT and C1HDT; Figure 4E); still, a large number of genes (e.g., Klf5, Tcl1, Klf2 and Nr0b1) showed a different pattern among differentiated samples C1T, C1MV, and C1HV, discriminating ESCs with different β-catenin doses (Figure 4E). These results support the hypothesis of a β-catenin-dependent effect on transcriptional changes.

A similar clustering across pluripotent samples was observed for general pluripotency markers (Figure 4F). In the majority of differentiated samples, Sox2 was downregulated while Utf1, Zfp281 and Lin28 were upregulated (Figure 4F), in accordance with previous reports (Zhang et al., 2016, Fidalgo et al., 2016, Luo et al., 2015, Betschinger et al., 2013). Under differentiated culture condition, Zfp281, Zic2 and Utf1 were downregulated in β-catenin induced cells (i.e., C1MV, C1HV, C1MDT and C1HDT) as compared to C1T ESCs (Figure 4F). Zfp281 is a Zinc finger transcription factor implicated in pluripotency (Brandenberger et al., 2004, Wang et al., 2008), and recently reported as a bidirectional regulator of the ESC-EpiSC transition in cooperation with Zic2, another zinc finger protein (Mayer et al., 2020). The undifferentiated embryonic cell transcription factor 1 (Utf1) is expressed in ESCs and plays an important role in the exit from pluripotency (Jia et al., 2012, Kooistra et al., 2009). The concomitant reduction of Zfp281, Zic2 and Urf1 in the comparison between C1T with both DMSO- and dox/TMP-treated samples suggests a global change in the chromatin organization of β-catenin induced ESCs en route to differentiation (Figure 4F). Finally, almost all the genes from this panel showed different behaviours in DMSO- (i.e., C1MV and C1HV; Figure 4F) *vs* dox/TMP-treated samples (i.e., C1MDT, C1HDT; Figure 4F), confirming that the extent of β-catenin induction affects cell identity.

Early post-implantation epiblast genes were mostly upregulated in primed ESCs compared to the pluripotent condition, with no evident differences across treatments in naïve ESCs (Figure 4G). The exception was Foxd3, which was downregulated in both naïve and primed β-catenin induced cells as compared to the controls C1 and C1T ESCs (Figure 4G). Interestingly, Dnmt3a and Dnmt3b showed a reduction in C1MV/C1HV and C1MDT/C1HDT samples as compared to the control C1T (Figure 4G); also, samples constantly exposed to dox/TMP (i.e., C1MDT and C1HDT) showed higher Dnmt3a expression than DMSO-treated ESCs (i.e., C1MV and C1HV; Figure 4G). Dnmt3a, b and Foxd3 are DNA and chromatin remodelling factors, respectively; Dnmt enzymes methylate genomic regions, whereas Foxd3 reduces active and enhances inactive histone marks by recruiting the Lysine-specific demethylase 1 (Lsd1) (Respuela et al., 2016). The reduced expression of those genes in β-catenin induced cells, including the pluripotent markers Utf1 discussed above, suggests that cells exposed to time/dose varying β-catenin levels present a differentially methylated DNA status during the exit from pluripotency.

We then screened for a large panel of lineage-priming factors. Ectoderm lineage markers showed a dose-dependent upregulation of related genes in pluripotent cells (compare C1, C1M and C1H; Figure 4H); following 2i/L withdrawal, the clustering resembled those of previous sets (Figures 4E-4G), with samples grouping for the duration of treatment (i.e., C1MV/C1HV and C1MDT/C1HDT grouping together; Figure 4H). Genes from this lineage had different expression across samples, making difficult to identify a clear pattern associated with β-catenin perturbations.

When looking at mesoderm markers (Figure 4I), differentiated samples clustered similarly to the previous data set. The first group of genes (i.e., Lhx1, Lefty1/2, Meox1, Hoxb1 and Bmp4) were mainly upregulated upon differentiation, whereas the second group (i.e., Nodal, Kdr, Mixl1, Gsc, Foxf1 and Zic1) got downregulated when exiting from pluripotency (Figure 4I). Although the pattern of individual genes was hard to interpret, we observed the behaviour of C1T ESCs was very different from all differentiated β-catenin induced samples, stressing the relevance of β-catenin for mesoderm specification (Lyashenko et al., 2011) and suggesting that its induction is diminishing mesoderm commitment.

The endoderm lineage was the most influenced by β-catenin perturbations: C1T cells were unable to induce the expression of endoderm-related genes (compare C1 and C1T; Figure 4J), whereas in all perturbed ESCs their expression increased over time. As previously observed (Figures 4E-4I), samples clustered together based on the duration of β-catenin induction rather than on the gene dose (i.e., C1MV/C1HV and C1MDT/C1HDT; Figure 4J). C1HDT cells showed the highest expression for the 50% of the endoderm-associated genes (namely, Cxcr4, Gata4 and Sox7) as compared to all other differentiated samples (i.e., C1T, C1MV, C1MDT and C1HV). These observations support previous knowledge about the β-catenin requirement for endoderm organization (Engert et al., 2013, Lyashenko et al., 2011).

In the analysis of the germ cell lineage markers, all genes showed a rather heterogeneous expression pattern across samples (Figure 4K). Pluripotent C1M and C1H clustered together and close to C1, and differentiated samples clustered based on the duration of β-catenin perturbation (i.e., C1MV/C1HV and C1MDT/C1HDT; Figure 4K).

Finally, when looking at trophectoderm markers (Figure 4L), clustering showed similarity of C1 and C1M, as for the ectoderm and endoderm lineages (Figures 4H and 4J, respectively). Of note, 90% of trophectoderm genes, with the exception of Cdx2, got downregulated during differentiation in all the conditions (Figure 4L). Eomes was recently reported to control the exit from pluripotency by acting on the chromatin status (Tosic et al., 2019); its behaviour in naïve C1M and C1H ESCs suggests a different chromatin conformation in pluripotent cells induced for β-catenin (Figure 4L).

Accounting for the fact that ectoderm is a default linage of the monolayer differentiation protocol we applied (Ying et al., 2003b), overall our sequencing results suggest that β-catenin induction in a knock-out background favours rescuing defects in differentiation towards endoderm more than mesoderm. Indeed, mesodermal genes were mostly downregulated when β-catenin was induced, whereas endodermal genes were all upregulated as compared to the control (Figures 4I and 4J). Moreover, we observed that lineage differentiation was influenced by the duration of protein induction rather than by its dose. Accordingly, there was a transition from mesoderm to endoderm following moderate but continuous β-catenin induction (compare C1MV and C1MDT in Figures 4A and 4B). Nevertheless, endoderm was an enriched gene ontology in all considered comparisons (Figures 4A-4D). Finally, the observed expression of pluripotency markers Zfp281, Zic2 and Utf1, the early post-implantation markers Dnmt3a-b and Foxd3 and the trophectoderm marker Eomes suggests a reorganization of the epigenome in naïve C1M and C1H ESCs and upon monolayer differentiation of C1MV, C1MDT, C1HV and C1HDT ESCs.

## Discussion

The role of the Wnt/β-catenin pathway as a pluripotency gatekeeper has been matter of many studies and debates (Sato et al., 2004, Ogawa et al., 2006, Hao et al., 2006, Singla et al., 2006, Anton et al., 2007, Takao et al., 2007, Kielman et al., 2002); while modulation of the canonical Wnt pathway has been extensively proved to be important for EpiSC *in vivo* derivation (Tsukiyama and Ohinata, 2014, Sugimoto et al., 2015), self-renewal (Sumi et al., 2013) and *in vitro* lineage differentiation (Liu et al., 2017, Osteil et al., 2019, Kurek et al., 2015), the relevance of β-catenin doses for the exit from pluripotency and for ESCs-EpiLCs direct transition has not been explored thoroughly.

In this work, we found that genetic β-catenin manipulation or chemical perturbation of the canonical Wnt pathway control ESC fate at the exit from pluripotency, providing new insights into the role of specific doses while confirm previous finding about the transcriptional role of β-catenin in pluripotency and early differentiation.

Using two different cellular models, we found that, upon FBS/L cultures, moderate β- catenin induction in differentiating β-catenin^-/-^ ESCs or moderate pre-activation of the Wnt/ β-catenin pathway in pluripotent wild type ESCs increase the efficiency of the ESCs-EpiLCs conversion. Pharmacological activation of the Wnt/β-catenin pathway in wild type ESCs gave different results as compared to genetic β-catenin induction in β-catenin^−/−^ ESCs (Figures 1, S1, 2 and S2). This observation could be explained by β-catenin-induced genetic variation reported in (Ortmann et al., 2020). Ortmann and colleagues demonstrated that β-catenin fluctuations in naïve pluripotent stem cells from different genetic backgrounds strongly influence how efficiently cells will differentiate (Ortmann et al., 2020). We believe that, by using β-catenin^−/−^ cells and inducing a β-catenin form which is insensitive to endogenous regulations, we abolished physiologic fluctuations and therefore mitigated the effect of genetic-variation on cell differentiation.

Simultaneous activation and inhibition of the Wnt/β-catenin pathway has been previously reported to maintain EpiSCs self-renewal (Kim et al., 2013): Kim and colleagues demonstrated that EpiSCs can be maintained in Chiron3µM/XAV2µM cultures with self-renewal regulated by both Axin2 and β-catenin. Our results suggest that the observation reported for EpiSCs (Kim et al., 2013) could also stand for the EpiLC derivation.

Overall, we confirmed the effect β-catenin has on preparing cells to appropriately respond to the differentiation stimuli previously reported (Ortmann et al., 2020), suggesting that both the duration and the dose of β-catenin induction control cell differentiation *in vitro*.

RNA sequencing performed in ESCs at the exit from the naïve ‘ground state’ of pluripotency (Kalkan et al., 2017) showed that, in β-catenin expressing cells (in particular C1MV), Dnmt3a and Dnmt3b had an expression pattern similar to the one observed in Rex1-high ESCs differentiated using a similar protocol (Kalkan et al., 2017), indicating that moderate β-catenin induction in naïve ESCs influences DNA methylation associated with the exit from pluripotency. β-catenin-dependent changes in DNA methylation have been previously reported in ESCs cultured for several passages in FBS/L (Theka et al., 2019). Theka and colleagues concluded that constant activation of the Wnt/β-catenin is necessary to guarantee adequate DNA methylation profiles. We also observed that persistent β-catenin induction (i.e., before and during differentiation, C1MDT and C1HDT; Figure 4G) partially restores the expression of Dnmt3a/b, which got downregulated following transient β-catenin induction (i.e., before differentiation, C1MV and C1HV; Figure 4G). Of note, we pre-cultured ESCs in 2i/L for 3 passages (1 week), a condition that strongly influences DNA methylation (Ficz et al., 2013, Habibi et al., 2013, Leitch et al., 2013). It would be interesting to assess the methylation state of specific genomic region, including imprinting control regions (ICRs), in response to dose- and time-varying β-catenin perturbations.

We observed a significant upregulation of endodermal genes in β-catenin induced cells, indicating a requirement of β-catenin for this specific fate. This phenotype was previously reported in the β-catenin null cell line generated by Lyashenko and colleagues (Lyashenko et al., 2011), where the defect in endoderm lineage differentiation was rescued by overexpressing both wild type or transcriptional incompetent β-catenin; in contrast, mesoderm and ectoderm induction seemed to not require β-catenin (Lyashenko et al., 2011). With our approach that enables dose- and time-controlled β-catenin induction, we also suggest that the ectoderm lineage is not affected by β-catenin loss. Different results were reported in (Tao et al., 2020), where β-catenin knockdown increased neural differentiation (Tao et al., 2020); given differences in the approaches used for β-catenin perturbations and culture conditions, additional studies (possibly also in other cell lines, and at the single-cell level) would be required for a direct comparison.

In the future, it will be of great interest to use our inducible system to interrogate the effect of a wider range of β-catenin doses and, possibly, temporal dynamics on stem- cell identity and to further investigate the role of the β-catenin transcriptional activity in pluripotent and differentiated cells of both murine and human origin (Antonella La Regina, 2021).

## Limitations of the study

We acknowledge the present study did not characterise the effect of high (i.e., above the wild-type levels) and/or dynamic β-catenin levels on cell decision making. Moreover, we did not consider comparing transcriptionally competent *vs* incompetent exogenous β-catenin; further studies uncoupling those two functions would be required and useful to fully unveil β-catenin-driven stem cell identity.

## STAR Methods

### Resource availability

#### Lead contact

Further information and requests can be addressed to Lucia Marucci or Elisa Pedone.

### Materials availability

This study did not generate new unique reagents.

### Experimental model and subject details Cell line derivation

C1 cell lines were previously derived in (Pedone et al., 2019) by a double lentiviral infection of β-catenin^-/-^ ESCs (Aulicino et al., 2020) with the EF1a-rtTA (Neomycin) plasmid followed by the pLVX_TrE3G-DDmCherryβ-catenin^S33Y^(Puromycin). Cells were selected with Neomycin after the first round and with Puromycin after the last infection.

ESCs were cultured on gelatin-coated dishes in Dulbecco’s modified Eagle’s medium (DMEM) supplemented with 15% fetal bovine serum (FBS), 1 x nonessential amino acids, 1 x GlutaMax, 1 x 2-mercaptoethanol, 1 x Penicillin-Streptomycin and 1000 U/mL LIF. To note, for the 2i/L culture, cells were kept for 3 passages (around 1 week) in serum-free NDiff227-based media supplemented with 1000 U/mL LIF, 3µM of the GSK-3α/β inhibitor Chiron-99021 and 1µM of the MEK inhibitor PD0325901.

### Method details

#### Epiblast-like cell (EpiLC) derivation

For EpiLC derivation *in vitro* we adapted the protocol for EpiSCs culture reported in (Kim et al., 2013), except that we employed N2B27 medium (Ying and Smith, 2003, Guo et al., 2009). Briefly, ESCs cultured in FBS/L or pre-cultured in 2i/L for 3 passages were seeded at the confluence of 1.5×10^4^ cells/cm^2^, on 10µg/mL Fibronectin-coated 12-well plates in NDiff227. According to the experiment in Figures 1D, 1E, S1G and S1H, cells were stimulated with DMSO, TMP10µM, dox10-100ng/mL, human ActivinA 10ng/mL and human FGF2 10ng/mL, whereas in Figures 2 and S2, cells were exposed to ActivinA 10ng/mL human FGF2 10ng/mL, Chiron1-3µM and the XAV939 2µM. Treatments were performed during 4 days with the media and drugs refreshed after the first 2 culture days. The concentration of ActivinA, human FGF2 and XAV939 were the same used in (Kim et al., 2013).

#### Monolayer differentiation

2i/L pre-cultured C1 ESCs were sorted based on β-catenin levels. Control C1, Middle (C1M) and High (C1H) expressing ESCs were plated at 1.5×10^4^ cells/cm^2^ on gelatin- coated 12-well plates in plain NDiff227 and stimulated with DMSO or TMP10µM±dox10-100ng/mL for 4 days with the media and drugs refreshed after the first 2 culture days (Figure 3A).

#### Drugs pre-treatment

Some experimental conditions required pre-treatment of cells. For β-catenin induction in Figures 1C-1E, S1G and S1H, C1 ESCs cultured in FBS/L or 2i/L were stimulated for 48 hrs with TMP10µM and dox10-100ng/mL before EpiSC differentiation, whereas for pre-activation of the canonical Wnt pathway in Figures 2A-2D, S2B-S2J and S2F- S2J, wild type ESCs were exposed for 48 hrs to Chiron1-3µM (Figures 2A-2D, S2B, S2C, S2F, S2G, S2I and S2J) or cultured for 3 passages in 2i/L (Figures 2A-2D, S2D and S2H-S2J), before the differentiation. To note, all experiments were performed with ESCs under heterogenous (i.e., FBS/L; Figures 1D, S1G, 2B-2D, S2A-S2C, S2E-S2G, S2I and S2J) or homogeneous (i.e., 2i/L; Figures 1E, S1H, 2B-2D, S2D and S2H-S2J) culture conditions. Transcriptional profiling was performed only with 2i/L pre-cultured ESCs (Figures 3, S3, 4 and S4).

#### Flow activated cell sorting (FACS)

2i/L pre-cultured ESCs were washed with sterile phosphate-buffered saline (PBS, Sigma), trypsinised for 2-3′ at room temperature and centrifuged at 1000 × g for 5′. Pelleted cells were resuspended in 500 μL of plain NDiff227 media supplemented with DAPI. The mCherry positive fraction was sorted from DAPI negative using the BD Influx high-speed 16-parameter fluorescence activated cell sorter.

#### qPCR

For quantitative PCR, the total RNA, extracted from cells using the PureLink RNA Mini Kit (Invitrogen), was retrotranscribed (Thermo Fischer, RevertAid Reverse Transcriptase) and the cDNA used as template for each qPCR reaction in a 15 μL reaction volume. iTaq Universal SYBR Green Supermix was used with the Qiagen Rotor-Gene System. To eliminate the contamination from genomic DNA, the RNeasy Plus Mini Kit (Qiagen) was used to purify the total RNA used for the RNA Sequencing.

#### QuantSeq 3’ RNA sequencing library preparation

Preparation of libraries was performed with a total of 100ng of RNA from each sample using QuantSeq 3’mRNA-Seq Library prep kit (Lexogen, Vienna, Austria) according to manufacturer’s instructions. Total RNA was quantified using the Qubit 2.0 fluorimetric Assay (Thermo Fisher Scientific). Libraries were prepared from 100ng of total RNA using the QuantSeq 3’ mRNA-Seq Library Prep Kit FWD for Illumina (Lexogen GmbH). Quality of libraries was assessed by using screen tape High sensitivity DNA D1000 (Agilent Technologies). Libraries were sequenced on a NovaSeq 6000 sequencing system using an S1, 100 cycles flow cell (Illumina Inc.).

Amplified fragmented cDNA of 300 bp in size were sequenced in single-end mode with a read length of 100 bp.

Illumina novaSeq base call (BCL) files are converted in fastq file through bcl2fastq.

#### QuantSeq 3’ RNA sequencing data processing and analysis

For analysis, sequence reads were trimmed using bbduk software (bbmap suite 37.31) to remove adapter sequences, poly-A tails and low-quality end bases (regions with average quality below 6). Alignment was performed with STAR 2.6.0a3 (Dobin et al., 2013) on mm10 reference assembly obtained from cellRanger website (https://support.10xgenomics.com/single-cell-gene-expression/software/release-notes/build#mm10_3.0.0; Ensembl assembly release 93). Expression levels of genes were determined with htseq-count (Anders et al., 2015) using Gencode/Ensembl gene model. We have filtered out all genes having < 1 cpm in less than n_min samples and Perc MM reads > 20% simultaneously. Differential expression analysis was performed using edgeR(Robinson et al., 2010), a statistical package based on generalized linear models, suitable for multifactorial experiments. The threshold for statistical significance chosen was False Discovery Rate (FDR) < 0.05 (GSE148879). The lists of differentially expressed genes (DEGs), for each comparison, with a threshold of logFC > 2 for the induced and logFC < -2 for the inhibited transcripts (Tables S2-S7) were used for the Functional Annotation analysis.

#### Weighted Gene Correlation Network Analysis (WGCNA)

Quant-seq 3’ mRNA data of 32 samples was used to construct a gene co-expression network by applying Weighted Gene Correlation Network Analysis (WGCNA)(Langfelder and Horvath, 2008) from the WGCNA package in the R statistical environment version 3.6. Briefly, we first computed the Pearson correlation coefficient among all pairs of expressed genes and then an appropriate value of the soft-thresholding power (β=6) giving a scale-free topology fitting index (R^2^) ≥ 0.85 was selected to build the weighted adjacency matrix. The weighted adjacency matrix was further transformed into a topological overlap matrix (TOM) (Yip and Horvath, 2007) and the resulting dissimilarity matrix used for hierarchical clustering. Gene modules were finally identified by cutting the hierarchical dendrogram with the dynamic tree cut algorithm from dynamicTreeCut package in R (Langfelder et al., 2008) statistical environment with standard parameters, except for cutHeight we set equal to 0.25 and deepSplit we set equal to 1. The value of deepSplit parameter was selected after performing a cluster stability analysis. Briefly, for each possible value of deepSplit parameter (i.e., 0, 1, 2, 3 or 4), modules were identified for both the full dataset and 50 resampled datasets. Then, the clustering solution obtained for the full dataset was compared with each resampled solution by mean of Adjusted Rand Index (ARI)(Hubert, 1985). The solution giving the highest average ARI was used for the clustering analysis as described above. Finally, to identify which clusters were correlated with β-catenin expression doses or differentiation time we correlated the first principal component of each gene module (i.e., the eigenmodule) with the traits of interest. The eigenmodule can be considered as a “signature” of the module gene expression. Modules correlated with the traits with a p-value < 0.01 were considered statistically significant and used for further analyses.

#### Functional Annotation Analysis

Differentially expressed genes (either logFC > 2 or logFC < -2) and module “hubs” having high module membership (also known as |KME| > 0.8) within the module were analysed for the enrichment in GO Biological Processes(Ashburner et al., 2000) and KEGG Pathways (Kanehisa and Goto, 2000) via the clusterProfiler package in R statistical environment (Yu et al., 2012). The threshold for statistical significance was FDR < 0.05, the top-ten BPs were represented as -log10 (FDR; Figures 3D, 3E, S3H- S3L, 4A-4D, S4A and S4B).

#### Statistical analysis

Differences between samples were analysed by two-tailed unpaired t-test and one-way ANOVA with Bonferroni’s multiple comparison test using GraphPad. A p-value lower than 0.05 was considered statistically significant.

Clustergram over heatmaps were generated using the clustergram function in Matlab that applies the Euclidean distance metric and average linkage. The data have been standardized across all samples for each gene and have 0 as mean and 1 as standard deviation.

## Data and code availability

- RNAseq raw data and analyses have been deposited on GEO: GSE148879. The GEO accession number is also listed in the key resources table.
- This paper did not report any original code.
- Additional information about this study is available from the lead contacts upon request.

## Acknowledgements

We thank Dr Andre Hermann and Dr Lorena Sueiro Ballesteros (Flow Cytometry Facility, University of Bristol), Dr Mark Jepson and Alan Leard (Wolfson Imaging Facility, University of Bristol) and the Next Generation Sequencing Core (TIGEM, Naples) for their support. This work was funded by Medical Research Council (grant MR/N021444/1) to L.M., by the Engineering and Physical Sciences Research Council (grants EP/R041695/1 and EP/S01876X/1 to L.M.), EC funding H2020 (FET OPEN 766840-COSY-BIO) to L.M., BrisSynBio, a BBSRC/EPSRC Synthetic Biology Research Centre (BB/L01386X/1) to L.M, STAR-University of Naples Federico II grant to G.G. and Fondazione Telethon grant to D.d.B.

## Authors Contributions

E.P. designed and performed experiments; M.F. and G.G. performed the WGNCA analysis; M.F. performed the GO; R.D.C. performed the Differential Expression analysis; ALR supported the experimental work; E.P. and L.M. analysed data; E.P., M.F., R.D.C. and L.M. wrote the paper; D.d.B. supervised the bioinformatics analysis; L.M. supervised the entire project.

## Declaration of interests

The authors declare that they have no competing interests.

## Notes

### Competing Interest Statement

The authors have declared no competing interest.

### Summary of Updates

In the new version of the manuscript, to confirm correct β-catenin we checked its levels before and/or during differentiation.

