## Supplemental Material for "β-catenin perturbations control differentiation programs in mouse embryonic stem cells"

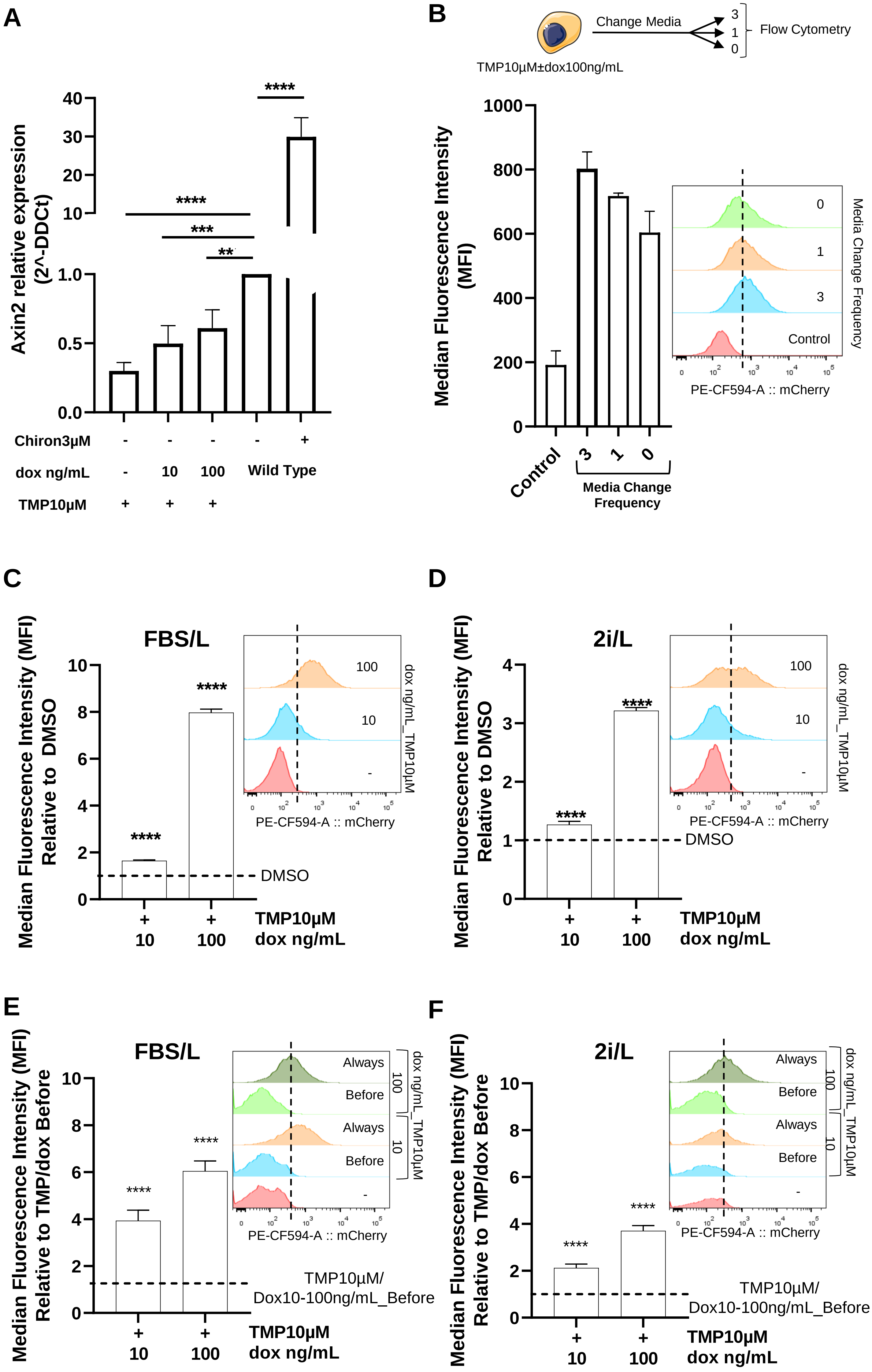
A
Axin2 relative expression (2^-DDCt)
40
30
20
10
1.0
0.5
0.0
****
****
***
**
Chiron3µM - - - - +
dox ng/mL - 10 100 Wild Type
TMP10µM + + +
B
TMP10µM±dox100ng/mL
Change Media
3
1
0
Flow Cytometry
Median Fluorescence Intensity (MFI)
1000
800
600
400
200
0
Control
3
1
0
Media Change Frequency
PE-CF594-A :: mCherry
Media Change Frequency
C
FBS/L
Median Fluorescence Intensity (MFI) Relative to DMSO
10
8
6
4
2
0
****
****
DMSO
100
10
-
dox ng/mL_TMP10µM
PE-CF594-A :: mCherry
+ + TMP10µM
10 100 dox ng/mL
D
2i/L
Median Fluorescence Intensity (MFI) Relative to DMSO
4
3
2
1
0
****
****
DMSO
100
10
-
dox ng/mL_TMP10µM
PE-CF594-A :: mCherry
+ + TMP10µM
10 100 dox ng/mL
E
FBS/L
Median Fluorescence Intensity (MFI) Relative to TMP/dox Before
10
8
6
4
2
0
****
****
Always
Before
Always
Before
-
100
10
dox ng/mL_TMP10µM
PE-CF594-A :: mCherry
TMP10µM/
Dox10-100ng/mL_Before
+ + TMP10µM
10 100 dox ng/mL
F
2i/L
Median Fluorescence Intensity (MFI) Relative to TMP/dox Before
10
8
6
4
2
0
****
****
Always
Before
Always
Before
-
100
10
dox ng/mL_TMP10µM
PE-CF594-A :: mCherry
TMP10µM/
Dox10-100ng/mL_Before
+ + TMP10µM
10 100 dox ng/mL

Figure S1

G

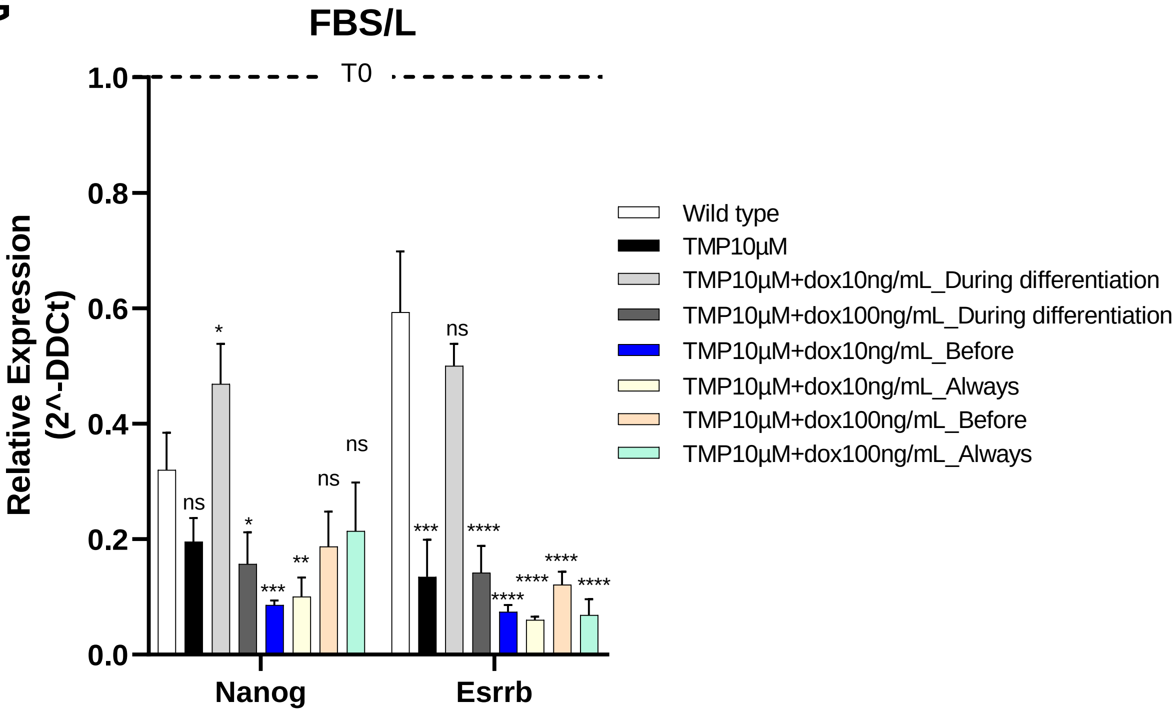
FBS/L
T0
Relative Expression
(2^-DDCt)
1.0
0.8
0.6
0.4
0.2
0.0
Nanog
Esrrb
Wild type
TMP10µM
TMP10µM+dox10ng/mL_During differentiation
TMP10µM+dox100ng/mL_During differentiation
TMP10µM+dox10ng/mL_Before
TMP10µM+dox10ng/mL_Always
TMP10µM+dox100ng/mL_Before
TMP10µM+dox100ng/mL_Always

H

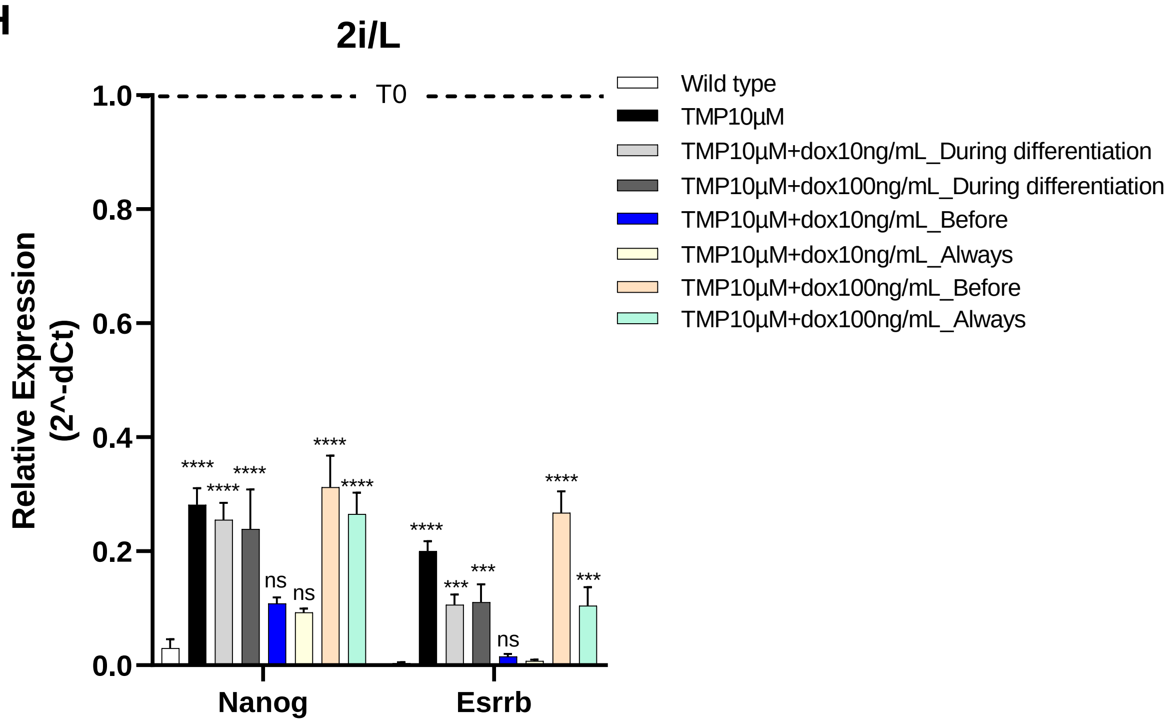
2i/L
T0
Relative Expression
(2^-dCt)
1.0
0.8
0.6
0.4
0.2
0.0
Nanog
Esrrb
Wild type
TMP10µM
TMP10µM+dox10ng/mL_During differentiation
TMP10µM+dox100ng/mL_During differentiation
TMP10µM+dox10ng/mL_Before
TMP10µM+dox10ng/mL_Always
TMP10µM+dox100ng/mL_Before
TMP10µM+dox100ng/mL_Always

Figure S1

**Figure S1. Characterization of β-catenin overexpressing C1 ESCs (Related to Figure 1).**

**A** Axin2 expression in C1 ESCs treated for 48 hrs with dox10-100ng/mL and/or TMP10μM, and wild type ESCs stimulated or not with Chiron3μM. **(B)** mCherry Median Fluorescence Intensity (MFI) of C1 ESCs cultured for 4 days in FBS/L supplemented with TMP10μM and dox10ng/mL. Media was changed and refreshed after 24 (3) and 48 (1) hrs in culture. Control cells were kept in the same media for the entire duration of the experiment without changing the media (0). **C-F** mCherry Median Fluorescence Intensity (MFI) of FBS/L **(C, E)** and 2i/L **(D, F)** C1 ESCs grown in pluripotent **(C, D)** and differentiating **(E, F)** culture conditions. Both pluripotent and differentiation media were supplemented with DMSO (i.e., time zero negative control or TMP10μM_dox10-100ng/mL_Before) or TMP10μM and dox10-100ng/mL (i.e., TMP10μM_dox10-100ng/mL_Always). Flow cytometry histograms are shown as inset. **G, H** Nanog and Esrrb expression in C1 ESCs cultured in FBS/L **(G)** or 2i/L **(H)** and differentiated for 4 days in NDiff227+ActivinA/FGF2 and different combination of DMSO, doxy and TMP. Data are represented as fold-change with respect to unstimulated wild type ESCs (A), DMSO-treated cells (C-F), or to the corresponding pluripotent condition (i.e., time zero before differentiation (T0) (G, H)); indicated with a dashed line. Data are means±SEM (n=3, A, C-H; n=2, B, biological replicates). p-values from two-tailed unpaired t test (A, C-F) and one-way ANOVA with Bonferroni's multiple comparison test (G, H) computed over the wild type (A, G, H) or DMSO-treated C1 (C-F) ESCs are shown, *p<0.05, **p<0.01, ***p<0.001, ****p<0.0001.

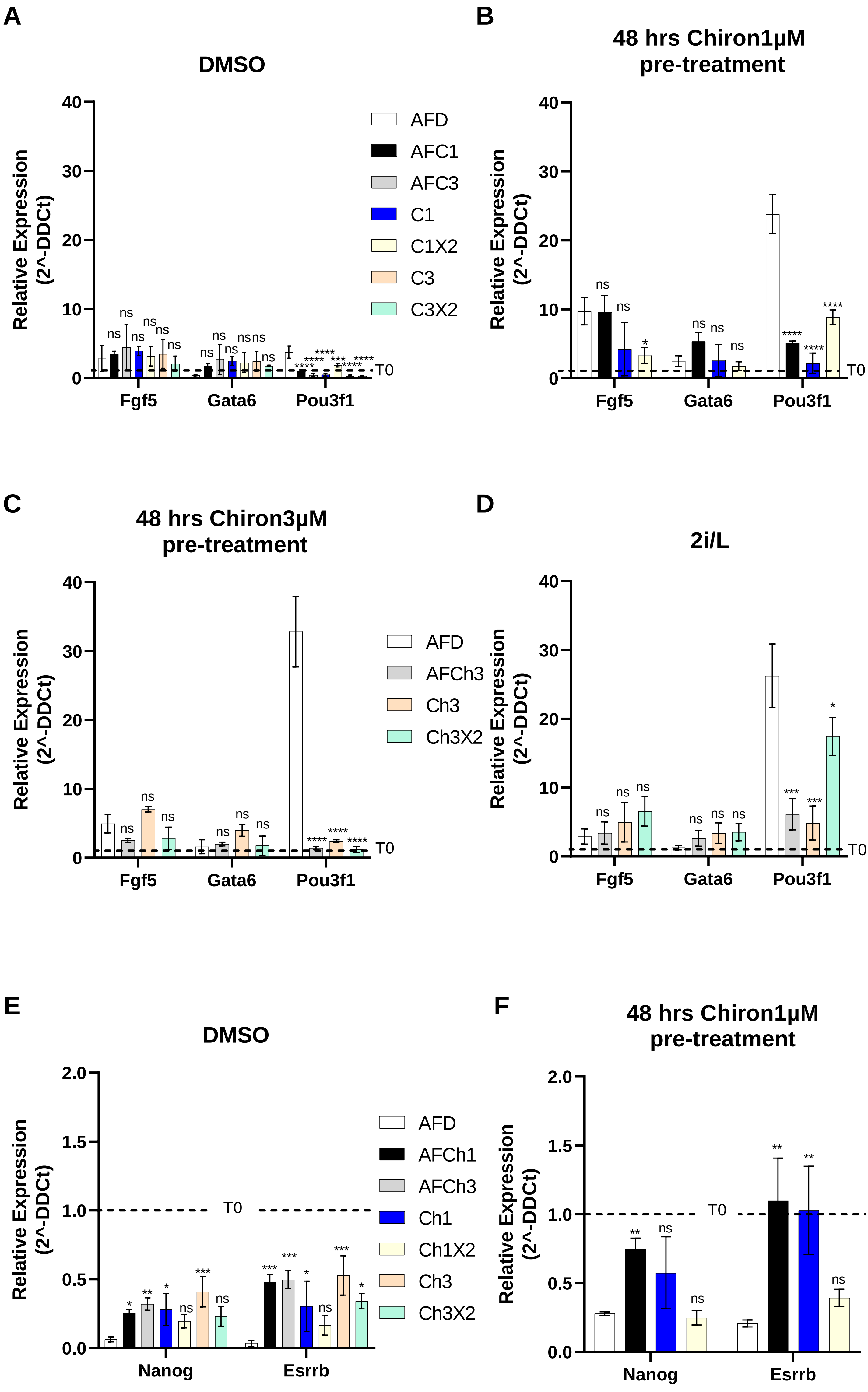

A
DMSO
B
48 hrs Chiron1µM pre-treatment
C
48 hrs Chiron3µM pre-treatment
D
2i/L
E
DMSO
F
48 hrs Chiron1µM pre-treatment
Relative Expression (2^-DDCt)
Fgf5
Gata6
Pou3f1
Nanog
Esrrb
T0
AFD
AFC1
AFC3
C1
C1X2
C3
C3X2
AFCh3
Ch3
Ch3X2
AFCh1
Ch1
Ch1X2


Figure S2

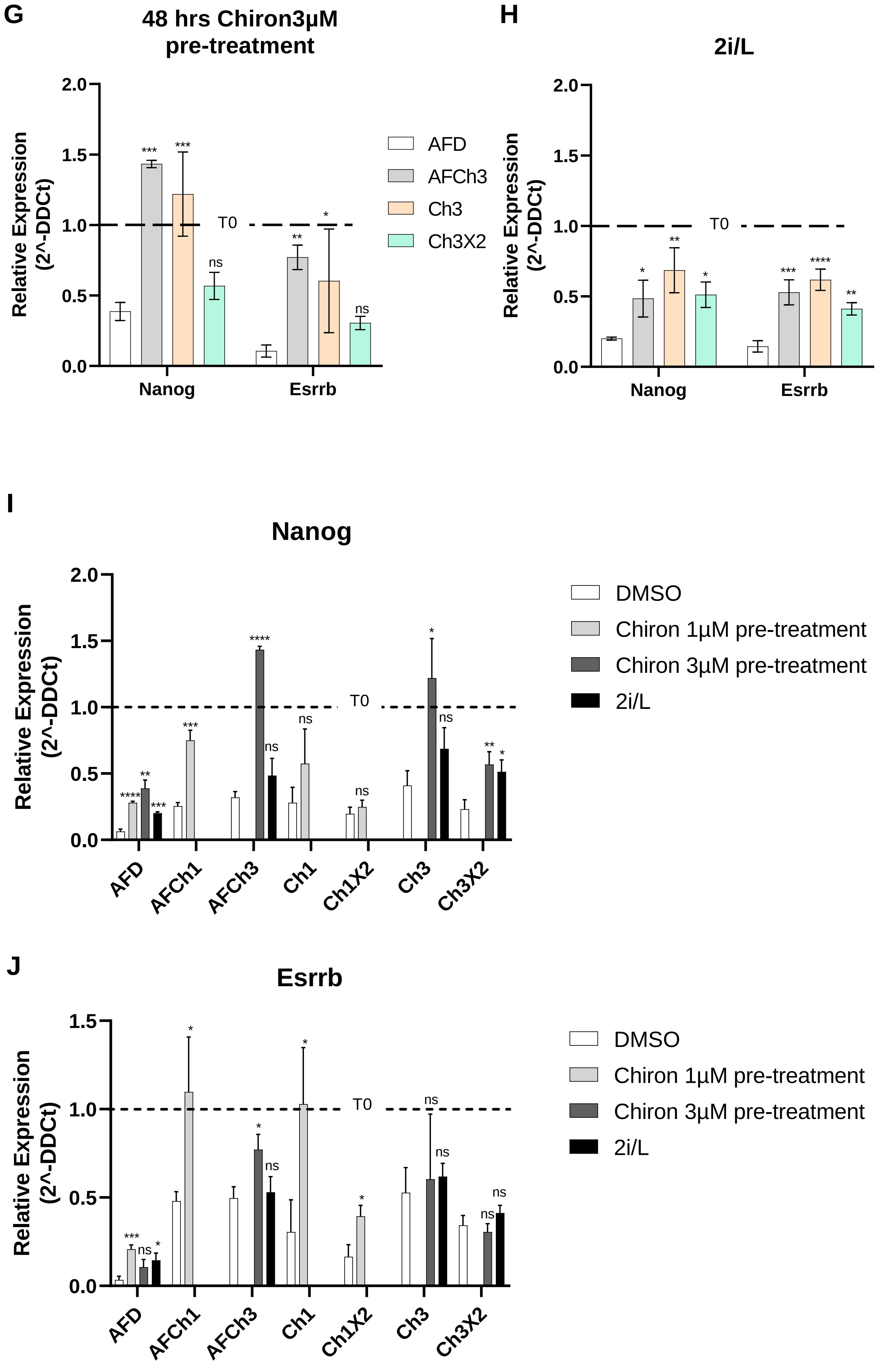
G
48 hrs Chiron3µM pre-treatment
Relative Expression (2^-DDCt)
T0
Nanog
Esrrb
AFD
AFCh3
Ch3
Ch3X2
H
2i/L
Relative Expression (2^-DDCt)
T0
Nanog
Esrrb
I
Nanog
Relative Expression (2^-DDCt)
T0
AFD
AFCh1
AFCh3
Ch1
Ch1X2
Ch3
Ch3X2
DMSO
Chiron 1µM pre-treatment
Chiron 3µM pre-treatment
2i/L
J
Esrrb
Relative Expression (2^-DDCt)
T0
AFD
AFCh1
AFCh3
Ch1
Ch1X2
Ch3
Ch3X2
DMSO
Chiron 1µM pre-treatment
Chiron 3µM pre-treatment
2i/L

Figure S2

**Figure S2. Characterization of wild type ESCs exposed to chemical perturbation of the Wnt/β-catenin pathway (Related to Figure 2).**

**A-D** Fgf5, Gata6 and Pou3f1 expression in DMSO **(A)**, Ch1µM **(B)**, Ch3µM **(C)** and 2i/L **(D)** pre-cultured wild type ESCs differentiated for 4 days in NDiff227 and the combination of drugs indicated in Figure 2. **E-H** Nanog and Esrrb expression in DMSO **(E)**, Ch1µM **(F)**, Ch3µM **(G)** and 2i/L **(H)** pre-cultured wild type ESCs differentiated for 4 days in NDiff227 and the combination of drugs indicated in Figure 2. **I, J** Nanog **(I)**, and Esrrb **(J)** expression in DMSO, Ch1µM, Ch3µM and 2i/L pre-cultured wild type ESCs differentiated for 4 days in NDiff227 and the combination of drugs indicated in Figure 2. Data are represented as fold-change with respect to the corresponding pluripotent condition (i.e., time zero before differentiation (T0), indicated with a dashed line). Data are means±SEM (n=3 biological replicates). p-values from one-way ANOVA with Bonferroni's multiple comparison test (A-H) and two-tailed unpaired t test (I, J) computed over the standard differentiation protocol based on ActivinA and FGF2 (i.e. AFD, A-H) or DMSO (I, J) are shown, *p<0.05, **p<0.01, ***p<0.001, ****p<0.0001.

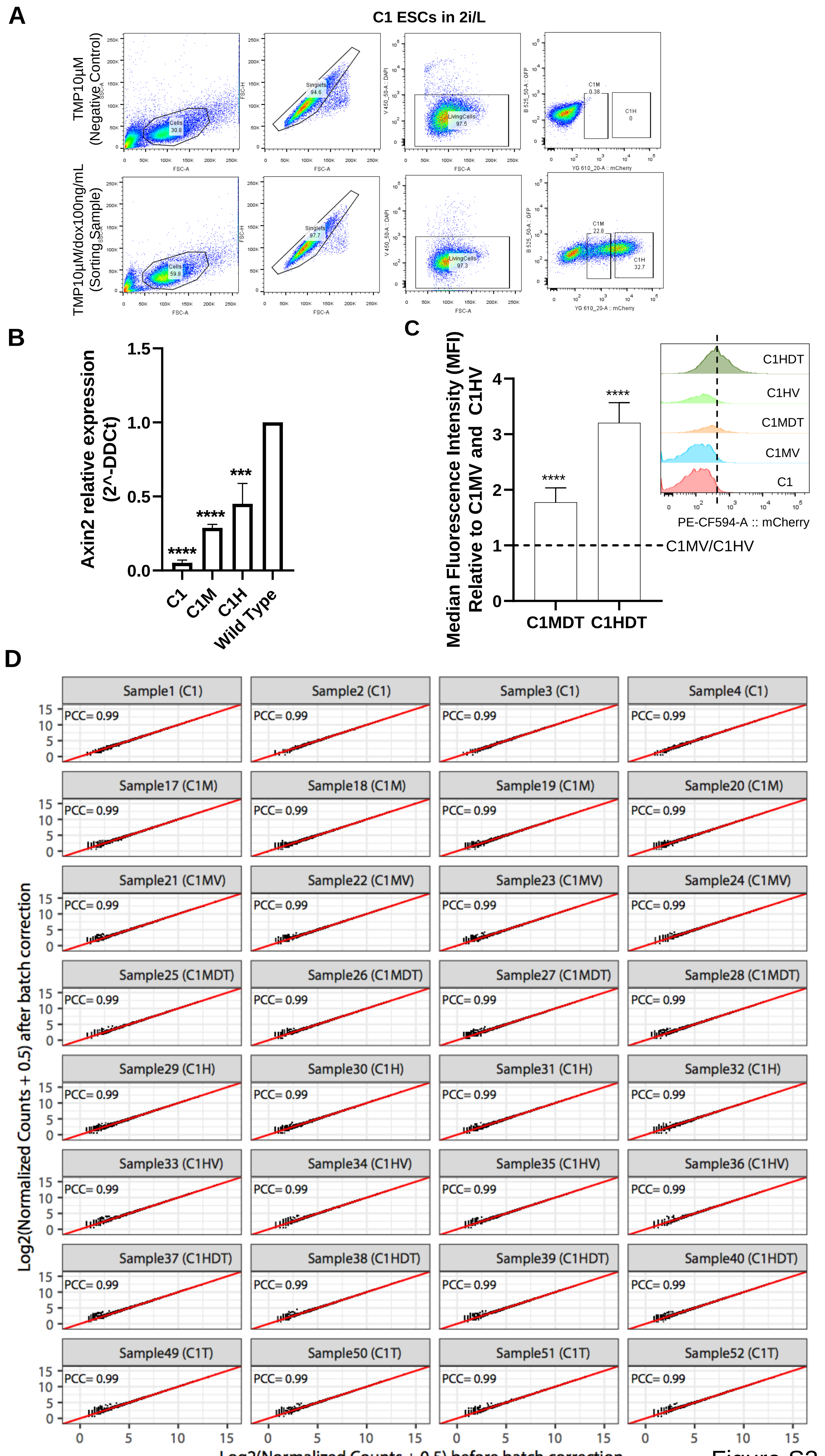
A
C1 ESCs in 2i/L
TMP10μM (Negative Control)
TMP10μM/dox100ng/mL (Sorting Sample)
Cells 30.8
Singlets 94.6
LivingCells 97.5
C1M 0.38
C1H 0
Cells 59.8
Singlets 97.7
LivingCells 97.3
C1M 22.8
C1H 32.7
FSC-A
YG 610_20-A :: mCherry
B
Axin2 relative expression (2^-DDCt)
C1
C1M
C1H
Wild Type
C
Median Fluorescence Intensity (MFI) Relative to C1MV and C1HV
C1MDT
C1HDT
C1MV/C1HV
C1HV
C1MV
C1
PE-CF594-A :: mCherry
D
Log2(Normalized Counts + 0.5) after batch correction
Log2(Normalized Counts + 0.5) before batch correction
PCC= 0.99
Sample1 (C1)
Sample2 (C1)
Sample3 (C1)
Sample4 (C1)
Sample17 (C1M)
Sample18 (C1M)
Sample19 (C1M)
Sample20 (C1M)
Sample21 (C1MV)
Sample22 (C1MV)
Sample23 (C1MV)
Sample24 (C1MV)
Sample25 (C1MDT)
Sample26 (C1MDT)
Sample27 (C1MDT)
Sample28 (C1MDT)
Sample29 (C1H)
Sample30 (C1H)
Sample31 (C1H)
Sample32 (C1H)
Sample33 (C1HV)
Sample34 (C1HV)
Sample35 (C1HV)
Sample36 (C1HV)
Sample37 (C1HDT)
Sample38 (C1HDT)
Sample39 (C1HDT)
Sample40 (C1HDT)
Sample49 (C1T)
Sample50 (C1T)
Sample51 (C1T)
Sample52 (C1T)

Figure S3

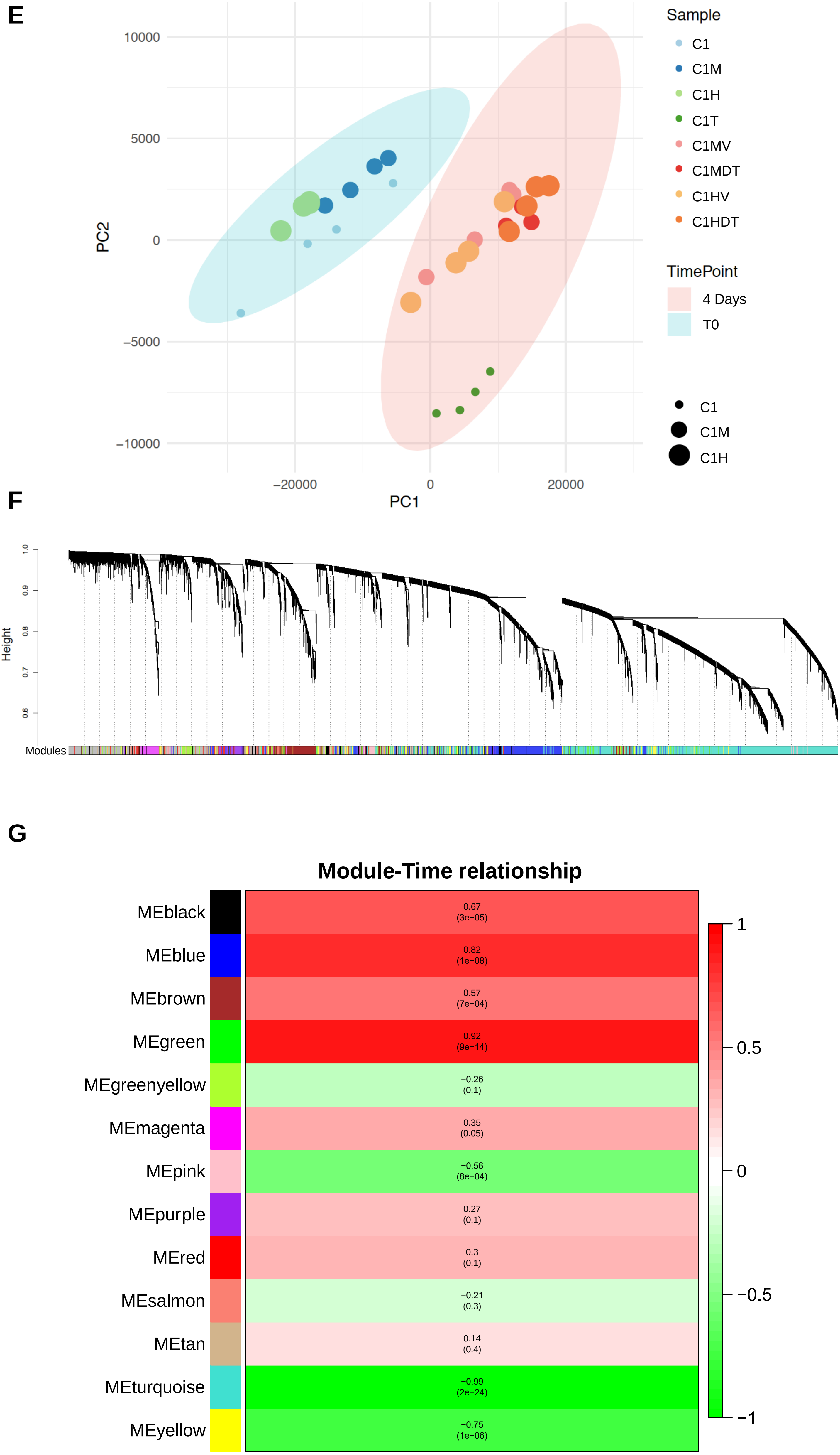
E
Sample
C1
C1M
C1H
C1T
C1MV
C1MDT
C1HV
C1HDT
TimePoint
4 Days
T0
C1
C1M
C1H
PC2
10000
5000
0
−5000
−10000
−20000
0
20000
PC1
F
Height
1.0
0.9
0.8
0.7
0.6
Modules
G
Module-Time relationship
MEblack
0.67 (3e−05)
MEblue
0.82 (1e−08)
MEbrown
0.57 (7e−04)
MEgreen
0.92 (9e−14)
MEgreenyellow
−0.26 (0.1)
MEmagenta
0.35 (0.05)
MEpink
−0.56 (8e−04)
MEpurple
0.27 (0.1)
MEred
0.3 (0.1)
MEsalmon
−0.21 (0.3)
MEtan
0.14 (0.4)
MEturquoise
−0.99 (2e−24)
MEyellow
−0.75 (1e−06)
1
0.5
0
−0.5
−1

Figure S3

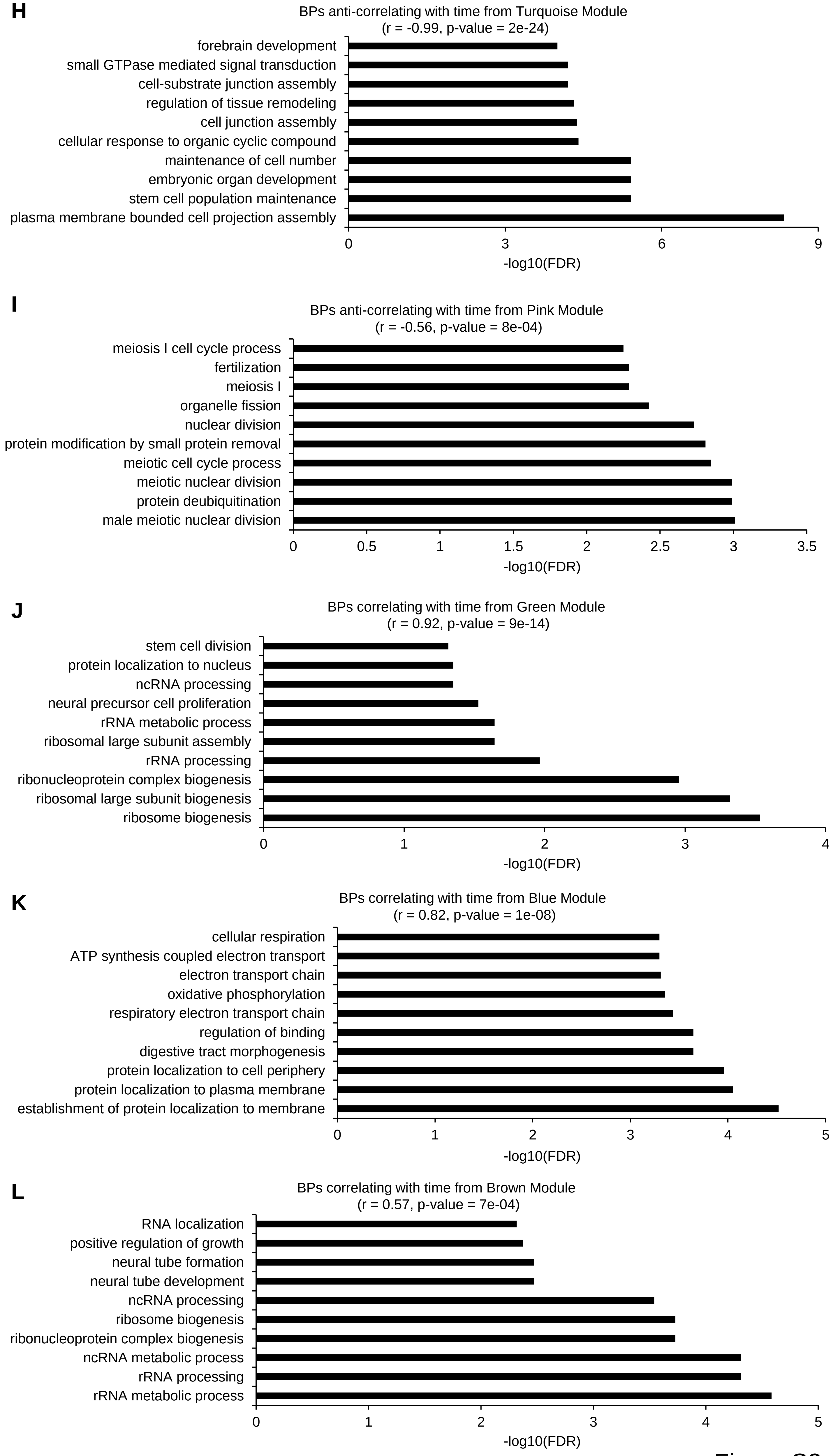
H
BPs anti-correlating with time from Turquoise Module
(r = -0.99, p-value = 2e-24)
forebrain development
small GTPase mediated signal transduction
cell-substrate junction assembly
regulation of tissue remodeling
cell junction assembly
cellular response to organic cyclic compound
maintenance of cell number
embryonic organ development
stem cell population maintenance
plasma membrane bounded cell projection assembly
0
3
6
9
-log10(FDR)
I
BPs anti-correlating with time from Pink Module
(r = -0.56, p-value = 8e-04)
meiosis I cell cycle process
fertilization
meiosis I
organelle fission
nuclear division
protein modification by small protein removal
meiotic cell cycle process
meiotic nuclear division
protein deubiquitination
male meiotic nuclear division
0
0.5
1
1.5
2
2.5
3
3.5
-log10(FDR)
J
BPs correlating with time from Green Module
(r = 0.92, p-value = 9e-14)
stem cell division
protein localization to nucleus
ncRNA processing
neural precursor cell proliferation
rRNA metabolic process
ribosomal large subunit assembly
rRNA processing
ribonucleoprotein complex biogenesis
ribosomal large subunit biogenesis
ribosome biogenesis
0
1
2
3
4
-log10(FDR)
K
BPs correlating with time from Blue Module
(r = 0.82, p-value = 1e-08)
cellular respiration
ATP synthesis coupled electron transport
electron transport chain
oxidative phosphorylation
respiratory electron transport chain
regulation of binding
digestive tract morphogenesis
protein localization to cell periphery
protein localization to plasma membrane
establishment of protein localization to membrane
0
1
2
3
4
5
-log10(FDR)
L
BPs correlating with time from Brown Module
(r = 0.57, p-value = 7e-04)
RNA localization
positive regulation of growth
neural tube formation
neural tube development
ncRNA processing
ribosome biogenesis
ribonucleoprotein complex biogenesis
ncRNA metabolic process
rRNA processing
rRNA metabolic process
0
1
2
3
4
5
-log10(FDR)

Figure S3

**Figure S3. FACS gating strategy and WGCNA of the genes correlating with time (Related to Figure 3).**

**A** Gating strategy used to sort C1M and C1H ESCs, following 48 hrs treatment with TMP10µM and dox100ng/mL. C1 ESCs treated with TMP10µM were used as negative control. **B** Axin2 expression in C1, C1M, C1H and wild type ESCs. **C** mCherry Median Fluorescence Intensity (MFI) of pluripotent C1MDT and C1HDT ESCs under differentiating culture conditions. Flow cytometry histogram is shown as inset. **D** Scatter plots of gene expression profiles, across all the 32 samples, before (x-axis) and after (y-axis) the batch correction. The Pearson correlation coefficient (PCC) is reported for each comparison. **E** Principal Component Analysis (PCA) of all samples; the average of replica is shown. **F** Clustering dendrogram of genes, with dissimilarity based on topological overlap, together with the assigned module colours**;** grey genes are unassigned to any module**. G** Eigenmodules correlating with time; the Pearson correlation coefficient (*r*) and relative p-value are shown. **H-L** Bar-chart of the top-ten enriched biological processes (BP) with FDR < 0.05 from genes belonging to the turquoise **(H)**, Pink **(I),** Green **(J)**, Blue **(K)** and Brown **(L)**, WGCNA modules.

Data are represented as fold-change with respect to wild type (B) or C1MV and C1HV (C, indicated with a dashed line) ESCs. p-values from two-tailed unpaired t test computed over the wild type (B) or C1MV and C1HV (C) ESCs are shown, *p<0.05, **p<0.01, ***p<0.001, ****p<0.0001.

**A**

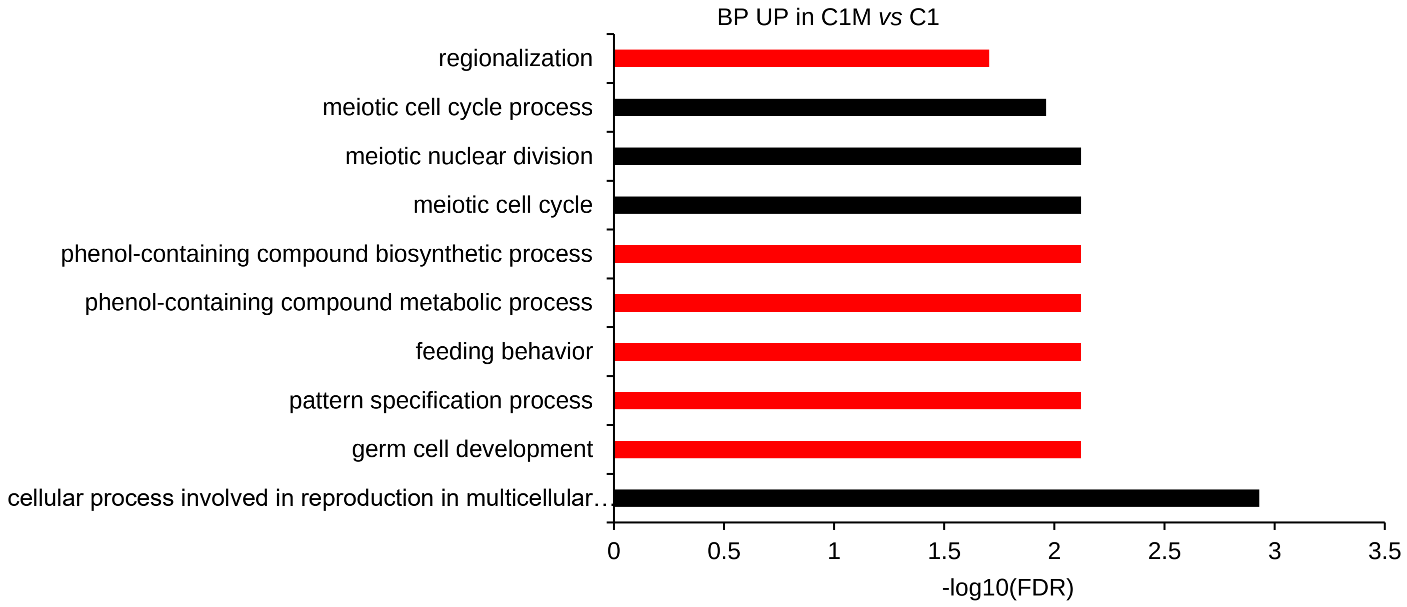

BP UP in C1M vs C1
regionalization
meiotic cell cycle process
meiotic nuclear division
meiotic cell cycle
phenol-containing compound biosynthetic process
phenol-containing compound metabolic process
feeding behavior
pattern specification process
germ cell development
cellular process involved in reproduction in multicellular…
0
0.5
1
1.5
2
2.5
3
3.5
-log10(FDR)


**B**

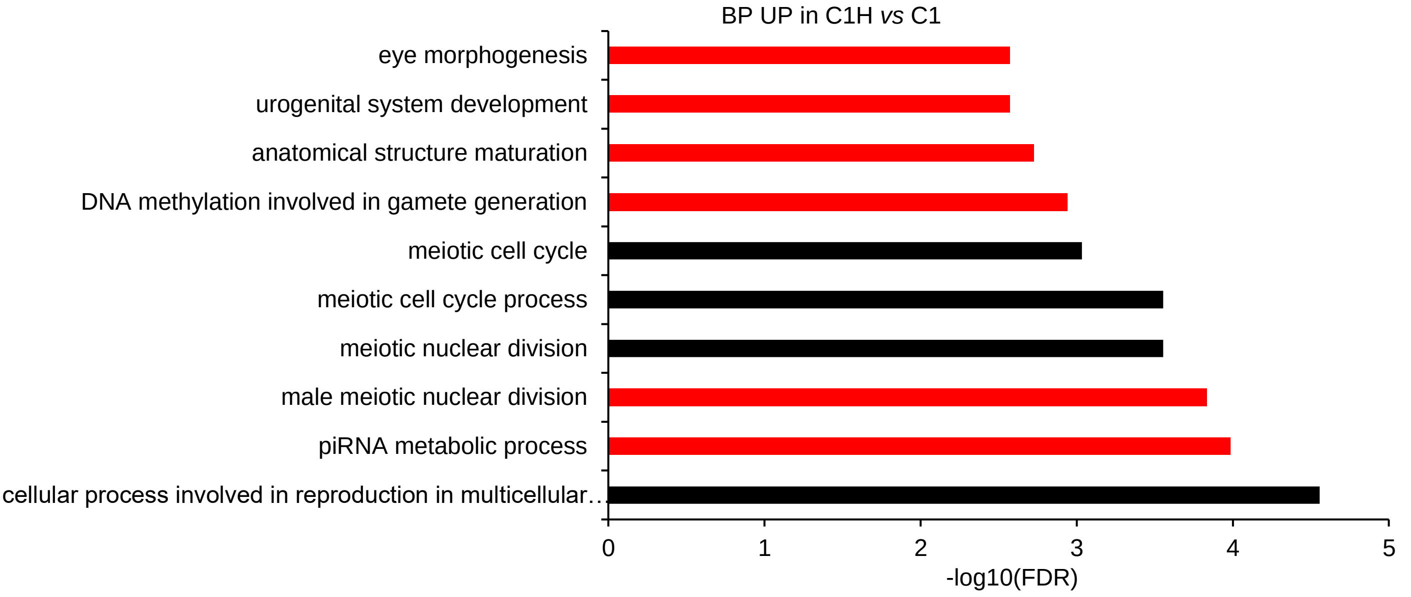

BP UP in C1H vs C1
eye morphogenesis
urogenital system development
anatomical structure maturation
DNA methylation involved in gamete generation
meiotic cell cycle
meiotic cell cycle process
meiotic nuclear division
male meiotic nuclear division
piRNA metabolic process
cellular process involved in reproduction in multicellular…
0
1
2
3
4
5
-log10(FDR)


Figure S4

**Figure S4. Gene ontology of the differential expressed genes in pluripotent ESCs (Related to Figure 4).**

**A, B** Bar-chart of the top-ten enriched biological processes (BP) with FDR < 0.05 from differentially expressed genes in in C1M **(A)** and C1H **(B)** compared to C1 ESCs. Black and grey bars represent upregulated and downregulated BPs, respectively. In red bars, the BPs exclusively enriched in the indicated condition.

**Table S1**

List of identified module genes correlating with time (Figure S3) or β-catenin doses (Figure 3) before and after the filtering for the |kME| ≥ 0.8; GO and pathway analysis.

**Table S2**

List of differentially expressed genes before and after the filtering for the FDR (< 0.05) and log fold change (-2 > log > 2); GO and pathway analysis in pluripotent C1M vs C1 ESCs.

**Table S3**

List of differentially expressed genes before and after the filtering for the FDR (< 0.05) and log fold change (-2 > log > 2); GO and pathway analysis in pluripotent C1H vs C1 ESCs.

**Table S4**

List of differentially expressed genes before and after the filtering for the FDR (< 0.05) and log fold change (-2 > log > 2); GO and pathway analysis in differentiated C1MV vs C1T ESCs.

**Table S5**

List of differentially expressed genes before and after the filtering for the FDR (< 0.05) and log fold change (-2 > log > 2); GO and pathway analysis in differentiated C1MDT vs C1T ESCs.

**Table S6**

List of differentially expressed genes before and after the filtering for the FDR (< 0.05) and log fold change (-2 > log > 2); GO and pathway analysis in differentiated C1HV vs C1T ESCs.

**Table S7**

List of differentially expressed genes before and after the filtering for the FDR (< 0.05) and log fold change (-2 > log > 2); GO and pathway analysis in differentiated C1HDT vs C1T ESCs.
